# BioMod: Biological Module Tokenization for Single-Cell Foundation Modeling

**DOI:** 10.64898/2026.10.12.758490

**Authors:** Lingbei Meng, Jianqing Zhu, Bowen Han, Yupeng Zang, Wenlin Li, Kewei Zong, Jian Zhong, Ying Yang, Jin Liu, Jiwei Jia, Ruoyu Sun, Guangjun Yu, Jiarui Han, Yan Chen, Lian Zhang, Jinchao Xu

**Affiliations:** Shenzhen Loop Area Institute; Shenzhen Research Institute of Big Data; The Chinese University of Hong Kong, Shenzhen; Shenzhen Raymind Biotechnology Co., Ltd.; National Health and Medical Big Data Research Institute (Shenzhen)

## Abstract

Single-cell foundation models typically represent cells as flat sequences of individual genes, requiring higher-order biological organization to be inferred implicitly from sparse expression profiles. Similar to the evolution from character-level to semantic tokenization in natural language processing,we hypothesize that cellular states can be more efficiently represented using biologically meaningful intermediate units. Here we present BioMod, a 50-million-parameter single-cell foundation model that introduces biological modules as representation units between genes and cells. BioMod constructs an overlapping vocabulary of 660 biological modules derived from curated pathways and data-driven gene programs, replacing thousands of gene-level inputs with compact module representations. Pretrained on approximately 73 million human cells using high-ratio module masking with an asymmetric decoder, BioMod achieves state-of-the-art performance across cell-type annotation, zero-shot batch integration, perturbation-response prediction and drug-response forecasting while maintaining a compact architecture that enables efficient inference on a single GPU. Module-level analyses further demonstrate that BioMod captures biologically meaningful cellular programs and preserves pathway-level organization.

## 1 Introduction

Single-cell RNA sequencing (scRNA-seq) provides a detailed view of cellular identities and states across tissues, development and disease [1]. Large transcriptomic atlases have enabled the development of single-cell foundation models (scFMs), which use self-supervised pretraining to learn representations for multiple downstream tasks [2]. Models including scGPT [3], Geneformer [4], scFoundation [5], UCE [6], scLong [7] and Cell2Sentence [8, 9] have been applied to cell-type annotation, data integration, perturbation-response prediction and drug-response modeling. Beyond architecture and training scale, the choice of representation unit remains an open question for single-cell foundation models.

Most existing scFMs use individual genes as the basic units of global encoding [3–5]. Gene-level tokens preserve individual expression measurements, but relationships among genes must be learned from sparse profiles. In language modeling, subword tokenization provides an intermediate granularity between individual characters and complete words. By analogy, biological modules might serve as representation units between individual genes and whole cells. Pathways, regulatory programs and molecular complexes group genes involved in coordinated cellular functions [10–12]. Unlike subwords, which partition a sequence without overlap, such modules overlap, are defined by prior knowledge or separate data analyses, and do not necessarily cover every gene. Representing these groups as module tokens could introduce biological structure into the input while shortening the sequence processed by the encoder. Whether such tokens retain the information needed for downstream tasks, and whether biological grouping contributes beyond sequence compression, must be tested empirically.

Gene sets have previously been incorporated into single-cell models in several ways. Map constrains latent dimensions to known gene programs for interpretable reference mapping [13], and TOSICA uses pathway-level tokens in a supervised transformer for cell-type annotation [14]. GeneCompass instead adds prior knowledge, including gene regulatory networks and co-expression relationships, to gene-level tokens during pretraining [15], and HEIMDALL has systematically examined tokenizer design in single-cell foundation models [16]. It remains unclear whether expression-conditioned module tokens can serve as the basic units of a self-supervised foundation model whose representations transfer across annotation, integration, perturbation and drug-response tasks.

To test this idea, we developed BioMod. BioMod represents each cell using a fixed vocabulary of 660 overlapping biological modules assembled from MSigDB Hallmark gene sets [11], Reactome pathways [12], Hotspot modules [17] and hdWGCNA co-expression modules [18]. This vocabulary combines curated biological knowledge with gene programs identified from single-cell expression data. Genes can belong to multiple modules, allowing their expression to contribute to different biological contexts. After identifier mapping, the modules cover 12,895 of the 27,874 genes in the model vocabulary; genes outside all modules do not enter module-token construction. Within each module, expression values scale Gene2Vec embeddings [19], and a learned attention pooler aggregates projected gene features into a cell-specific token. The global encoder therefore processes module tokens rather than individual gene tokens. During pretraining, 75% of the module tokens are removed from the encoder input, and an asymmetric decoder reconstructs gene-expression values within the masked modules. This design pairs a short encoder sequence with reconstruction supervision at gene resolution.

BioMod contains approximately 50 million parameters and was pretrained on approximately 73 million human cells across 25 organ/tissue categories. We evaluated the model on cell-type annotation, zero-shot batch integration, genetic perturbation prediction and drug-response prediction, with comparisons against foundation models containing up to one billion parametersand task-specific methods. To assess the contribution of biological gene grouping, we also pretrained a shuffled-module model with the same module count, module-size distribution, architecture and training protocol. This comparison tests gene-to-module assignments within a fixed module-token architecture; comparisons with larger models assess the performance achievable at the reported parameter count. Module-family ablations, marker validation and pre-encoder gene-set sensitivity and post-encoder module masking were used to examine the representations supporting cell-type classification.

BioMod achieved the highest macro-F1 on the four annotation datasets with tabulated baseline comparisons, the highest Pearson correlation in three of four perturbation-prediction settings(although its MSE ranking varied across settings), and the highest scores on both drug-response benchmarks among the compared models. Shuffling gene-to-module assignments reduced overall integration scores in all four evaluation conditions, annotation macro-F1 on all seven datasets and performance on both drug-response tasks, whereas perturbation-prediction effects varied across settings and metrics. In cell-type classification, highly ranked modules contained canonical markers, and masking these modules caused a greater loss of accuracy than masking low-ranked or random modules. Thus, a compact model built on biological modules supports diverse single-cell tasks. The controlled comparisons indicate that biological gene grouping contributes to performance in several tasks, and module-level analyses link classifier decisions to defined gene programs.

## 2 Results

### 2.1 Overview of BioMod

BioMod uses a fixed vocabulary of 660 overlapping biological modules (Fig. 1c). This module set comprises 50 MSigDB Hallmark gene sets [11], 491 Reactome pathways [12], 73 Hotspot modules [17], and 46 hdWGCNA modules [18] (Fig. 1c). Hallmark and Reactome provide curated pathway definitions, whereas Hotspot and hdWGCNA contribute gene programs derived from single-cell expression data. Modules overlap by design, so a gene can contribute to more than one module when it occurs in several gene sets.

**Figure 1.**
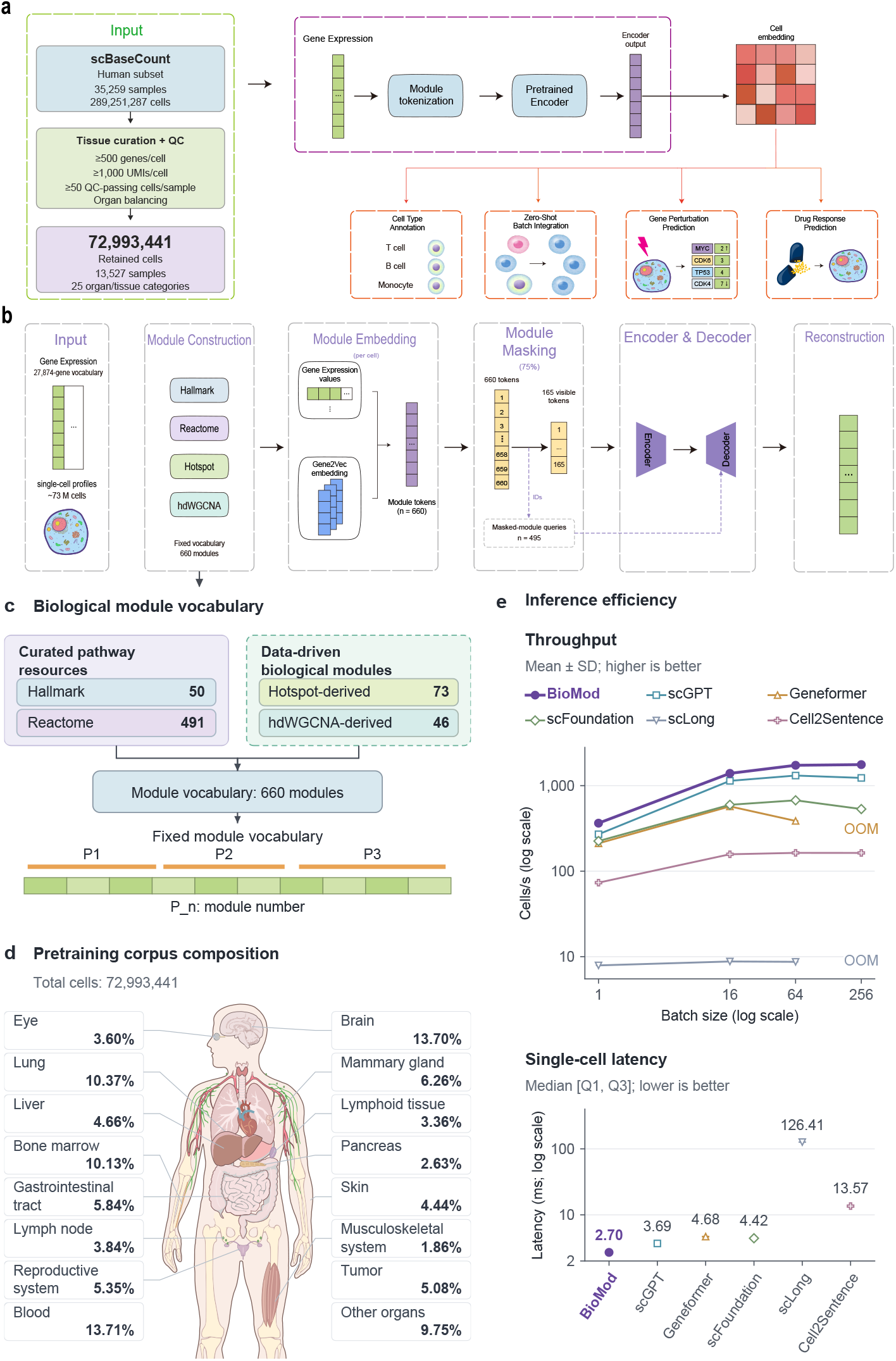
Overview of BioMod: data curation, biological module pretraining, downstream applications and inference efficiency. **(a)** Overall workflow. Curation, quality control and organ balancing of the human scBaseCount subset yielded 72,993,441 cells from 13,527 samples across 25 organ/tissue categories. Gene-expression profiles are converted into module tokens and processed by a pretrained encoder to produce cell embeddings for cell-type annotation, zero-shot batch integration, gene-perturbation prediction and drug-response prediction. **(b)** Cell-specific module tokenization and masked pretraining. Expression values and Gene2Vec gene embeddings are combined to construct 660 module tokens per cell. At a 75% masking ratio, 165 visible tokens enter the encoder, while 495 masked-module queries enter the decoder for gene-resolution reconstruction of expression in the masked modules. **(c)** Fixed biological module vocabulary, comprising 50 Hallmark gene sets [11], 491 Reactome pathways [12], 73 Hotspot-derived modules [17] and 46 hdWGCNA-derived modules [18]. *P*_*n*_ denotes a module’s position in the vocabulary. **(d)** Organ/tissue composition of the pretraining corpus. The anatomical schematic displays selected categories, with the remaining categories grouped as “Other organs”. **(e)** Representation-forward efficiency of BioMod and five comparator models on an NVIDIA H100 80-GB GPU. Upper, throughput at batch sizes of 1, 16, 64 and 256 cells (mean ± sample standard deviation across five technical repeats; logarithmic axes). Lower, batch-size-1 forward latency (median and interquartile range; logarithmic latency axis). The BioMod curve used the guarded pool-affine implementation and 512 cells per repeat; comparators used 10,000 cells per repeat from the same selected cohort. Cross-model differences are therefore descriptive. OOM denotes failed batch-size-256 runs of Geneformer and scLong.

For each cell, BioMod constructs one token for each of the 660 modules before global sequence modeling. Within module *p*_*i*_, the expression value *x*_*g*_ of each constituent gene scales its Gene2Vec embedding **e**_*g*_, and learned weights combine the resulting gene features into one module representation. This differs from reducing a pathway to a fixed enrichment score because the representation depends on the gene-expression pattern of the current cell. The global encoder consequently receives 660 module tokens rather than the more than 2,000 individual gene tokens used by representative gene-level models [3, 5].

The module sequence is processed by a TSSA encoder [20]. During pretraining, 75% of the module tokens are removed from the encoder input, leaving 165 visible tokens per cell. An asymmetric decoder uses the visible module states to reconstruct the expression values of individual genes in the masked modules. The encoder therefore operates on a shorter sequence, while the pretraining loss remains defined at gene resolution for genes in the masked modules.

We pretrained BioMod on approximately 73 million human cells across 25 organ/tissue categories, curated from scBaseCount [21] (Fig. 1d; Methods). The complete model has approximately 50 million parameters. For downstream evaluation, the backbone was fully fine-tuned for cell-type annotation, used as a frozen feature extractor for zero-shot integration and drug-response prediction, and connected to a module-to-gene back-projection for perturbation-response prediction. We first report task-specific results and then compare the model sizes and performance of BioMod with those of the baseline foundation models.

Because module tokenization shortens the sequence processed by the encoder, we also measured inference cost directly (Fig. 1e; Supplementary Tables S14 and S15). We timed the representation forward pass, from learned input processing through the backbone to cell-level pooling, on a single NVIDIA H100 80-GB GPU with BF16 autocast, using Zheng68K cells and excluding CPU preprocessing, host-to-device transfer and unused decoders or output heads. We compared BioMod with scGPT, Geneformer, scFoundation, scLong and Cell2Sentence. BioMod had the highest throughput at every batch size tested. At batch size 1 it processed 364.1 ± 4.7 cells per second, compared with 270.0 ± 1.1 for scGPT, 224.7±1.5 for scFoundation, 211.3±0.9 for Geneformer, 73.3±0.3 for Cell2Sentence and 7.9 for scLong. Its median single-cell latency was 2.70 ms, compared with 3.69 ms for scGPT, 4.42 ms for scFoundation, 4.68 ms for Geneformer, 13.57 ms for Cell2Sentence and 126.41 ms for scLong, with narrow interquartile ranges for all models. The advantage widened with batching: throughput of BioMod rose to 1,391 cells per second at batch size 16 and plateaued at about 1,730–1,761 cells per second at batch sizes 64 and 256, where it was about 1.4 times that of scGPT (1,231 at batch size 256), 3.3 times that of scFoundation (532) and 10.8 times that of Cell2Sentence (163). At batch size 64, it exceeded Geneformer by 4.5 times and scLong by about 200 times. Throughput of Geneformer, scFoundation and scGPT declined at larger batch sizes, and Geneformer and scLong ran out of GPU memory at batch size 256, whereas BioMod completed all settings. These differences are consistent with the shorter module-token sequence (660 tokens, compared with more than 2,000 gene tokens in gene-level models) and the linear-complexity TSSA encoder. They are descriptive rather than matched estimates, since BioMod was timed on 512 cells per repeat and the comparators on 10,000, each model kept its own input adapter and sequence length, and we did not separate the contributions of tokenization, attention mechanism and implementation.

### 2.2 Cell-type annotation

Accurate cell-type identification is a foundational task in single-cell biology, essential for constructing cell atlases, characterizing tissue composition, and understanding disease heterogeneity [2]. Cell-type annotation requires assigning each cell to a discrete identity from its transcriptomic profile [3, 4]. To evaluate how well module-level representations encode cell-identity information, we benchmarked BioMod on seven datasets spanning diverse tissues and annotation granularities: Kim Lung, Immune, Liver [22], MS, hPancreas [14], Zheng68K [23], and Myeloid [24] (workflow in Fig. 2a; Supplementary Tables S1 and S2), comparing against scGPT [3], Geneformer [4], scFoundation [5], and Cell2Sentence (the 410-million-parameter C2S-Scale model) [9] where corresponding baseline results were available.

**Figure 2.**
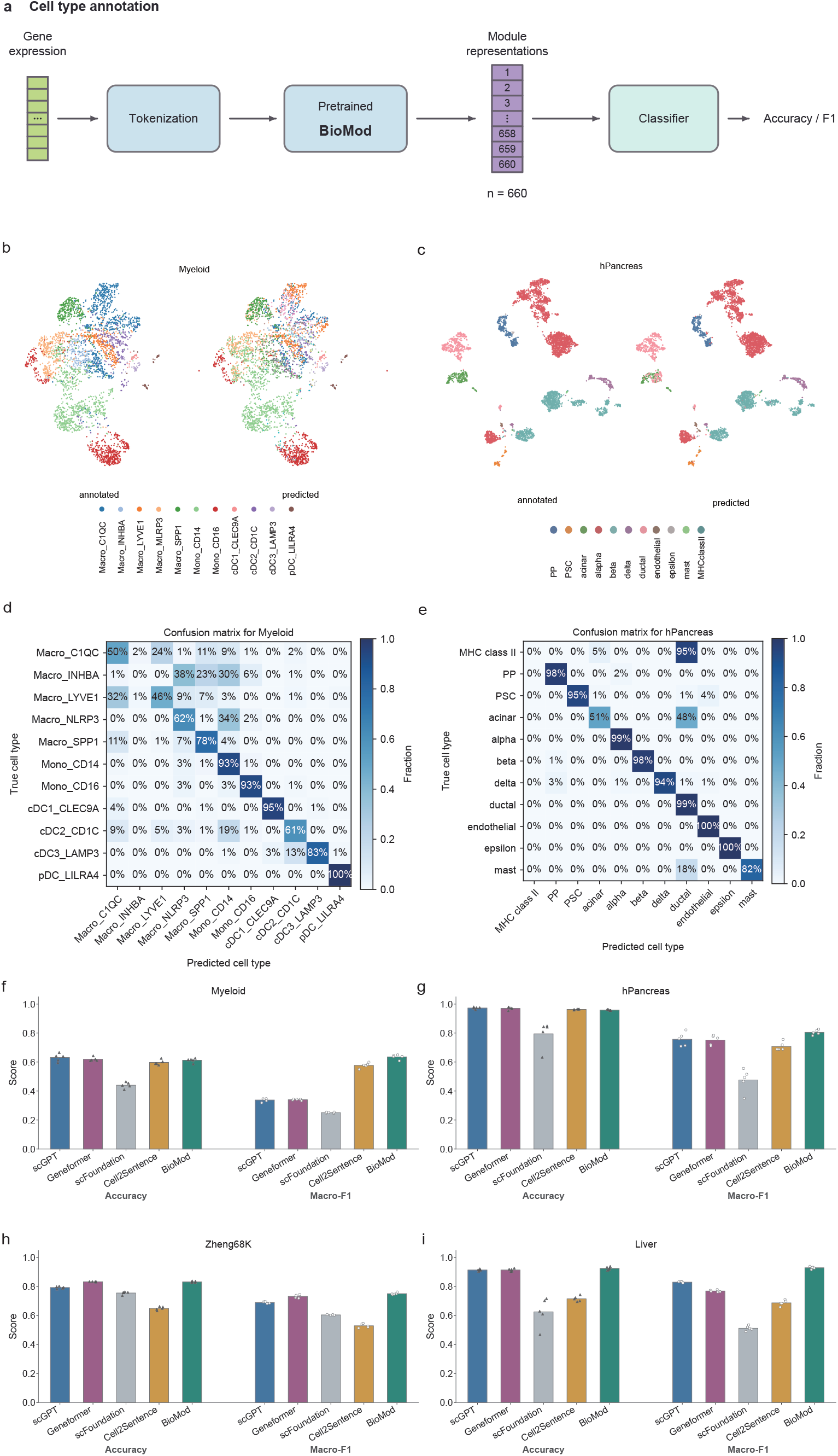
BioMod annotates cell types across diverse single-cell datasets. **(a)** Schematic of the cell-type annotation workflow. **(b)** Two-dimensional visualization of the Myeloid [24] dataset. **(c)** Two-dimensional visualization of the hPancreas [14] dataset. **(d)** Normalized confusion matrix for Myeloid. **(e)** Normalized confusion matrix for hPancreas. **(f–i)** Benchmark comparisons on Myeloid, hPancreas, Zheng68K and Liver, respectively, showing accuracy and macro-F1 for scGPT, Geneformer, scFoundation, Cell2Sentence and BioMod.

The performance of BioMod varied across the annotation benchmarks, with direct base-line comparisons shown for Myeloid, hPancreas, Zheng68K and Liver (Fig. 2f–i). Across five independent runs, it reached mean accuracy and macro-F1 of 0.913 and 0.918 on Liver, respectively. Its macro-F1 exceeded those of scGPT (0.831) and Geneformer (0.770), while its accuracy was slightly below their shared value of 0.915. On Zheng68K, BioMod achieved the highest accuracy (0.839) and macro-F1 (0.752) among the compared methods, exceeding Geneformer (0.834/0.733). The corresponding accuracy and macro-F1 values were 0.986/0.962 on Kim Lung, 0.948/0.930 on Immune, and 0.956/0.797 on hPancreas. Performance on MS and Myeloid was more modest (0.793/0.679 and 0.615/0.645). On Myeloid, BioMod achieved the highest macro-F1 despite lower accuracy than scGPT and Gene-former. Across the four datasets with tabulated baseline comparisons, BioMod achieved the highest macro-F1, whereas its accuracy ranking varied by dataset (Supplementary Tables S1 and S2).

On hPancreas, predicted labels recovered most major pancreatic populations, including endocrine, stromal, endothelial, and several exocrine cell types, as shown by the two-dimensional visualization and confusion matrix (Fig. 2c, e). However, MHC class II-positive and acinar cells showed greater cross-class confusion. On the more challenging Myeloid dataset, which spans closely related myeloid subtypes from multiple cancer types, BioMod resolved monocyte and dendritic-cell populations more consistently than several closely related macrophage subtypes, with the annotated and predicted cell-type structures broadly aligned in embedding space (Fig. 2b, d).

Overall, representations learned through module-token pretraining retained enough information to classify most major cell populations, whereas performance was lower for several closely related subtypes. These experiments do not indicate whether the subtype-level errors arise from module coverage, class imbalance, training-set size or other properties of the individual datasets.

### 2.3 Zero-shot batch integration

Single-cell datasets generated across different laboratories, sequencing platforms, and experimental batches frequently exhibit systematic technical variations, known as batch effects, that can obscure genuine biological signals and confound downstream analyses [25]. To assess whether the frozen module representation generalizes across technical conditions, we evaluated BioMod on zero-shot batch integration, a task that tests whether cell embeddings can separate biological variation from systematic technical artifacts [25] without access to batch labels or domain-adaptation training. We compared BioMod against Gene-former [4], scGPT [3], scFoundation [5], and UCE [6] on three heterogeneous atlases: the hPNC dataset [25] (16,382 cells, 6 batches, 14 subtypes), BMMC [25] (90,261 cells, 12 batches), and a human lung atlas [26] (32,493 cells, 14 patient batches, 10 lineages), using the scIB metric suite [25] (Methods).

Across three datasets (four evaluation conditions), BioMod showed dataset-dependent biological conservation and batch mixing over five independent runs (seeds 0–4; Fig. 3; Supplementary Table S3). Because malignant cells can exhibit strong patient-specific transcriptional programs that may be confounded with patient-level batch structure [26], we evaluated the lung atlas both with and without malignant cells. Excluding malignant cells increased the overall score from avg_score_ = 0.781 ± 0.006 to 0.851 ± 0.012 (Fig. 3d, e), with a corresponding rise in the biological-conservation score from avg_bio_ = 0.647 ± 0.010 to 0.761 ± 0.020, while batch mixing remained high (avg_batch_ = 0.982 ± 0.000 and 0.986 ± 0.000, respectively). This suggests that tumor-specific heterogeneity may obscure the non-malignant lineage organization captured by the frozen representation. On hPNC and BMMC, BioMod achieved overall scores of 0.665 ± 0.003 and 0.662 ± 0.006, respectively, with batch-mixing scores of 0.901 ± 0.006 and 0.907 ± 0.002 (Fig. 3b, c). Compared with Cell2Sentence [9], BioMod achieved higher overall scores on Kim Lung (0.781 versus 0.766), hPNC (0.665 versus 0.645), and Kim Lung without malignant cells (0.851 versus 0.786), but a lower score on BMMC (0.662 versus 0.677).

**Figure 3.**
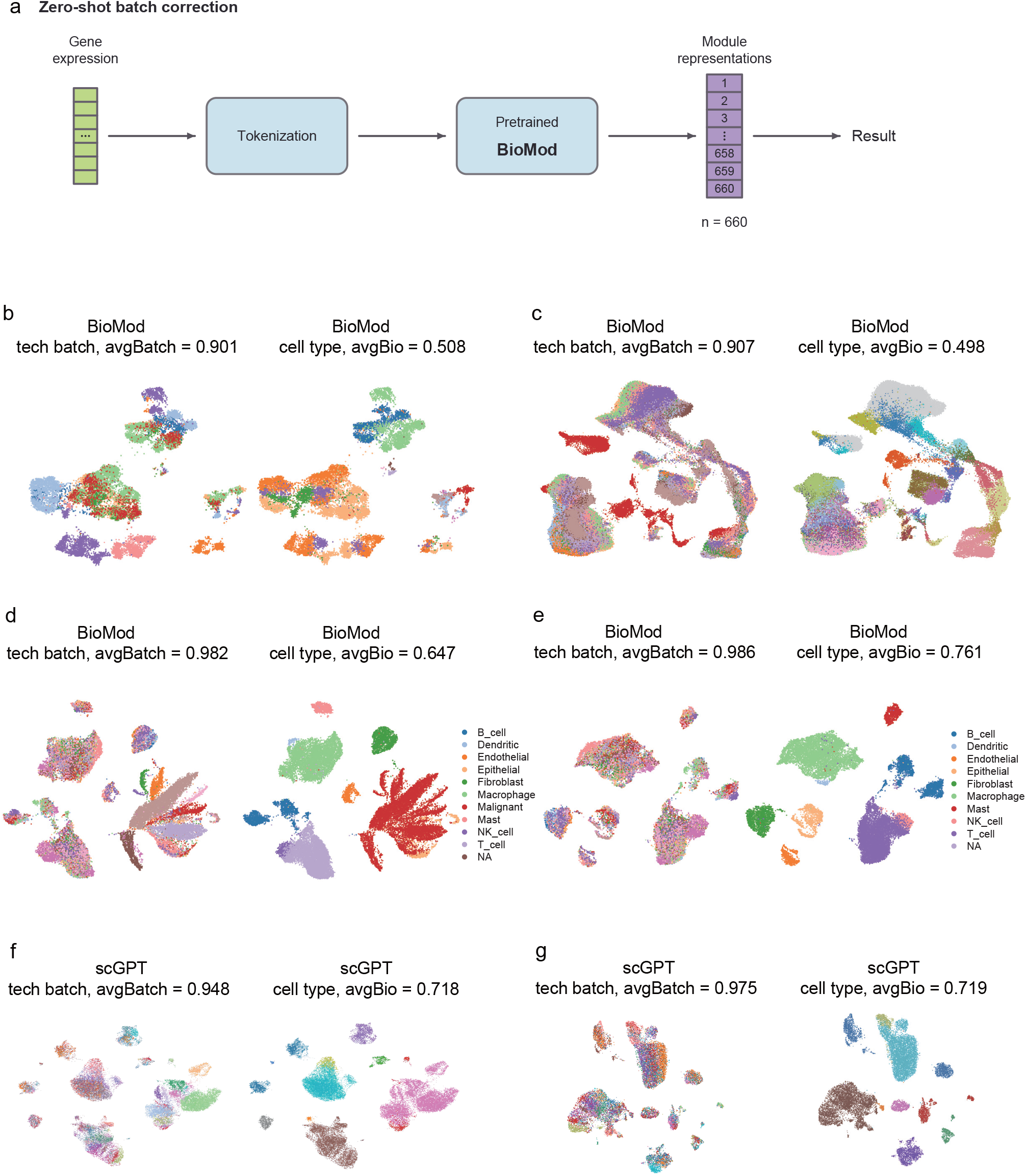
Zero-shot integration with BioMod preserves biological structure across heterogeneous datasets. **(a)** Schematic of the zero-shot inference workflow: gene expression profiles are mapped to the frozen BioMod encoder and aggregated into cell embeddings without task-specific fine-tuning. **(b–e)** Quantitative annotations report five-run means for avg_bio_ as the biological-conservation score, avg_batch_ as the batch-mixing score and avg_score_ as the weighted overall score. **(b)** UMAP on the hPNC dataset [25], colored by batch and cell type (avg_bio_ = 0.508, avg_batch_ = 0.901, avg_score_ = 0.665). **(c)** UMAP on BMMC [25] (avg_bio_ = 0.498, avg_batch_ = 0.907, avg_score_ = 0.662). **(d)** UMAP on Kim Lung [26] with malignant cells retained (avg_bio_ = 0.647, avg_batch_ = 0.982, avg_score_ = 0.781). **(e)** UMAP on Kim Lung after removing malignant cells (avg_bio_ = 0.761, avg_batch_ = 0.986, avg_score_ = 0.851). **(f, g)** Corresponding scGPT [3] zero-shot embeddings on Kim Lung with and without malignant cells, showing more fragmented geometry.

UMAP visualizations support these quantitative results. In the lung setting, BioMod produces a globally organized manifold in which immune, stromal, epithelial, and malignant populations are resolved, with non-malignant lineages remaining clearly structured after malignant cells are excluded (Fig. 3d, e). By comparison, scGPT yields a more fragmented geometry, with isolated clusters and weaker global relationships among non-malignant lineages (Fig. 3f, g). On hPNC and BMMC, BioMod similarly reduces batch-driven separation while preserving cell-type boundaries (Fig. 3b, c). Thus, the frozen module representation achieves competitive zero-shot integration without an explicit domain-alignment objective, although specialized methods with explicit batch supervision [27, 28] remain important comparators for dedicated integration tasks.

In the tested datasets, the frozen representation preserved cell-type structure while mixing technical batches, without batch labels or task-specific adaptation. Because the evaluation concerns the complete pretrained model, it does not separate the contributions of module tokenization, encoder architecture and pretraining objective, and it is not direct evidence that module grouping alone reduces gene-level dropout noise.

### 2.4 Perturbation-response prediction

Understanding how genetic perturbations reshape the transcriptome is central to functional genomics and therapeutic target discovery, yet experimentally profiling all possible perturbation conditions remains infeasible [29, 30]. We used perturbation-response prediction to test whether representations learned through module-token pretraining can support a task requiring gene-level outputs. BioMod encodes the pre-perturbation profile as module representations, which are projected back to genes and combined with GEARS-derived perturbation embeddings to predict post-perturbation expression [30] (Fig. 4a; Methods). We evaluated the Norman dataset [29] (91,205 cells, 5,045 genes and 236 perturbation conditions) under four generalization settings (Seen 0/1, Seen 0/2, Seen 1/2 and Seen 2/2), using Pearson correlation and MSE on the top 20 differentially expressed genes. Comparisons included Geneformer [4], scGPT [3], scFoundation [5], UCE [6], scLong [7], and the task-specific GEARS model [30] (Fig. 4c; Supplementary Table S5).

**Figure 4.**
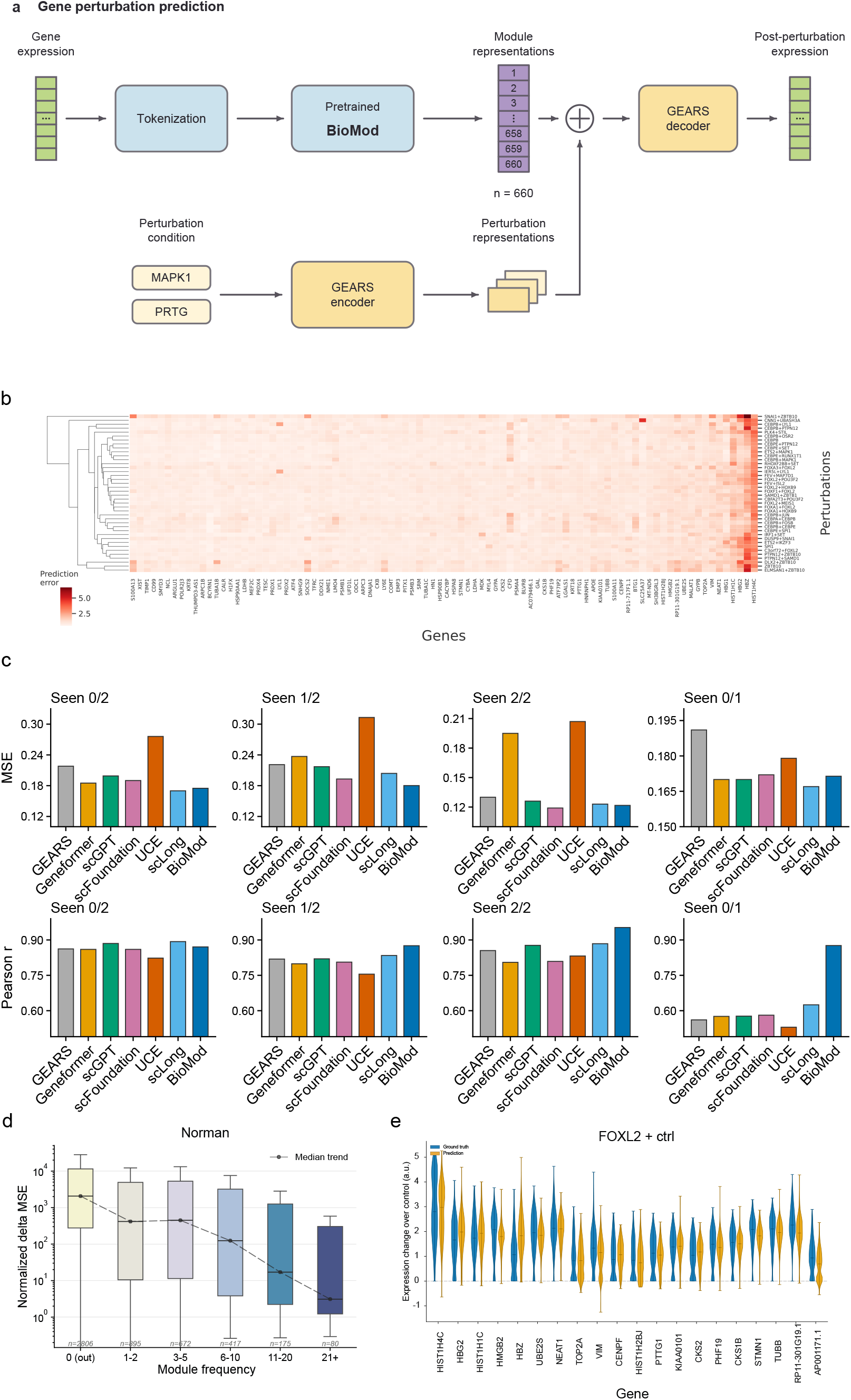
BioMod predicts transcriptional responses to genetic perturbations. **(a)** Schematic of the perturbation-response prediction framework. **(b)** Heatmap of prediction errors across perturbation conditions (rows) and genes (columns) in the Norman dataset. **(c)** Benchmark comparison across four generalization settings (Seen 0/1, 0/2, 1/2, 2/2), showing MSE and Pearson correlation on the top 20 differentially expressed genes. **(d)** Association between the number of modules containing each gene and prediction error. **(e)** Violin plots comparing observed and predicted expression changes for the *FOXL2* +ctrl perturbation.

BioMod achieved the highest Pearson correlation in three of the four generalization settings, whereas its MSE ranking varied by setting. In Seen 0/1, it obtained *r* = 0.876 and MSE = 0.171: Pearson correlation exceeded that of the strongest baseline, scLong (*r* = 0.625), but MSE was higher than scLong’s 0.167. In Seen 1/2, BioMod ranked first in both metrics (*r* = 0.876, MSE = 0.180). In Seen 2/2, it achieved the highest Pearson correlation (*r* = 0.953), while its MSE of 0.122 was slightly higher than the best value of 0.119 from scFoundation. In Seen 0/2, BioMod obtained *r* = 0.871 and MSE = 0.175, compared with *r* = 0.893 and MSE = 0.170 for scLong. Overall, BioMod achieved the lowest MSE in one setting and the highest Pearson correlation in three, so its advantage was more consistent in correlation than in squared error. These comparisons assess the complete prediction pipeline. The shuffled-module control below separately examines whether biological gene grouping contributes to this task.

Fig. 4b summarizes BioMod’s mean absolute prediction errors across the 90 genes and 40 perturbation conditions with the largest errors. Hierarchical clustering showed that some conditions sharing a perturbed gene had similar error profiles; for example, CEBPB+LYL1 and PTPN12+CEBPB were grouped together. Errors were concentrated in a subset of genes and perturbation conditions; this descriptive analysis does not identify the regulatory properties responsible for those differences.

Because prediction errors varied across genes, we examined their association with each gene’s coverage within the set of 660 modules. In the Norman dataset, normalized delta MSE decreased as the number of modules containing a gene increased (Fig. 4d). This association is consistent with well-covered genes appearing in more module contexts, but it does not show that module overlap lowered the error, because coverage may covary with expression level, annotation depth, gene function or other gene properties. For the representative *FOXL2* +ctrl perturbation, the predicted distributions followed the observed expression changes for most of the displayed differentially expressed genes, with larger mismatches for a subset of targets (Fig. 4e).

### 2.5 Cross-modal drug-response prediction

Predicting how cancer cells respond to therapeutic compounds is a central problem in precision oncology, as patients with the same histological diagnosis can exhibit different drug sensitivities [31, 32]. We used this task to test whether representations learned by the module-token model transfer to a setting that combines intracellular transcriptomic state with the structure of an external compound [32, 33]. Cell-line expression profiles were represented as 660 module tokens and encoded by the frozen BioMod backbone. The resulting cell representation was combined with drug molecular features to predict the natural-log-transformed half-maximal inhibitory concentration (ln(IC_50_)) (Fig. 5a; Methods).

**Figure 5.**
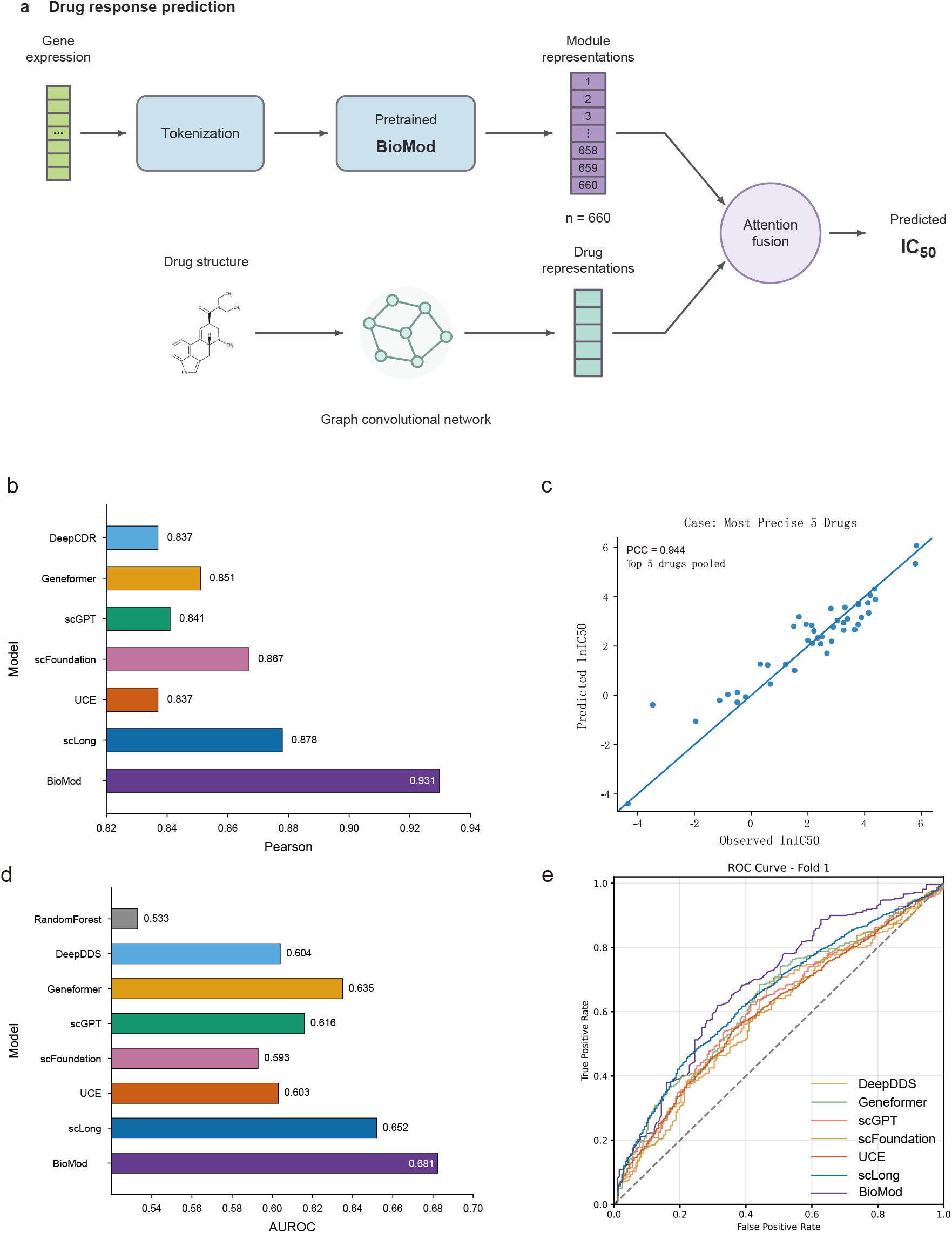
BioMod predicts single-drug and drug-combination responses. **(a)** Schematic of the drug-response prediction framework. Gene-expression profiles are represented as 660 module tokens and encoded by the frozen BioMod backbone. Drug molecular structures are encoded by a graph convolutional network. The resulting cell-state and drug representations are combined by an attention-fusion module to predict ln(IC_50_). **(b)** Single-drug response prediction on the CCLE–GDSC benchmark test set. Bars show Pearson correlation between predicted and observed ln(IC_50_) values for DeepCDR, Geneformer, scGPT, scFoundation, UCE, scLong and BioMod. The BioMod bar reports the five-seed mean (*r* = 0.931). **(c)** Predicted versus observed ln(IC_50_) values for the five most accurately predicted drugs pooled across cell lines. Each point represents a drug–cell-line pair. The panel reports *PCC* = 0.944. This selected subset is illustrative; overall benchmark performance is reported in panel b and Supplementary Table S6. **(d)** Drug-combination response prediction benchmark. Bars show AUROC for RandomForest, DeepDDS, Geneformer, scGPT, scFoundation, UCE, scLong and BioMod. The BioMod bar reports the five-seed mean (AUROC = 0.681). **(e)** Receiver operating characteristic curves for the drug-combination prediction task. These curves were computed on the same drug-combination test set as panel (d). The dashed diagonal denotes the random-classifier reference.

We evaluated BioMod on the CCLE–GDSC benchmark [31, 34, 35] (107,446 matched records, of which 94,314 were retained after cancer-type filtering; a validation set was used for checkpoint selection, and approximately 18,863 records were reserved as the test set for final evaluation under a record-level split; Methods), comparing against five foundation models: scLong [7], Geneformer [4], scGPT [3], scFoundation [5], and UCE [6], as well as the task-specific model DeepCDR [34]. BioMod achieved the highest reported Pearson correlation among the compared models (Fig. 5b, c; Supplementary Table S6), with a five-seed mean of 0.931 ± 0.003, compared with 0.878 for scLong, 0.867 for scFoundation, 0.851 for Geneformer, 0.841 for scGPT, and 0.837 for DeepCDR.

We next extended the evaluation to drug-combination synergy prediction, in which the output is a binary label indicating whether a drug pair exhibits synergy. We trained on a large-scale oncology screening dataset (*>* 12,000 examples) and tested on an independent AstraZeneca screen (668 samples) [36]. BioMod outperformed all reported baselines in AU-ROC (Fig. 5d, e; Supplementary Table S7), attaining a five-seed mean of 0.681 ± 0.004 versus 0.652 for scLong, 0.635 for Geneformer, 0.616 for scGPT and 0.604 for the task-specific model DeepDDS. Together with the single-drug result, this indicates that embeddings from the module-token encoder carry information predictive of both single-compound response and drug-combination synergy under the evaluated protocols. These comparisons do not isolate the effect of the module-token design from differences in pretraining data, backbone architecture or downstream fusion models.

### 2.6 A 50-million-parameter module model performs comparably to larger foundation models

A 660-token module design achieved strong downstream performance with substantially fewer parameters. BioMod contains approximately 50 million parameters, compared with 1 billion for scLong. BioMod was pretrained on approximately 73 million cells, whereas scLong was pretrained on approximately 48 million cells. Under the reported benchmarks, BioMod outperformed scLong on five of eight perturbation-prediction metrics and on both drug-response tasks (*r* = 0.931 versus 0.878 for single-drug response and AUROC = 0.681 versus 0.652 for drug-combination response). BioMod also contains approximately one eighth as many parameters as the reported 410-million-parameter Cell2Sentence model and achieved higher macro-F1 on all four cell-annotation datasets with tabulated comparisons, together with higher accuracy on Zheng68K, Liver and Myeloid but lower accuracy on hPancreas (0.956 versus 0.963) (Supplementary Tables S2 and S4). Thus, a 50-million-parameter module-token model can reach the reported downstream performance. These comparisons do not show that the module-token design alone accounts for the differences between models, because the models differ in pretraining-corpus size, architecture, objectives and downstream adaptation. The module-shuffling experiments below address the narrower question of whether biological gene-to-module assignments matter when module count, module sizes, model architecture and training protocol are held fixed.

### 2.7 Ablation studies

We next tested two properties of the module design. First, we asked whether downstream performance changes when biological gene-to-module assignments are disrupted while module count, module-size distribution, architecture and training protocol are held fixed. Second, we examined classification sensitivity to the gene sets associated with the four module sources. These experiments test the gene-to-module organization within BioMod; they do not provide a matched comparison between module-token and gene-token architectures.

#### Shuffled-module control

To separate the effect of biological module organization from that of gene grouping alone, we constructed a shuffled-module control in which genes were randomly permuted and reassigned to modules according to the original module sizes. This preserved the number of modules, the module-size distribution, the model architecture and the training protocol, but disrupted functional relationships among genes within each module. The shuffled-module model was pretrained from scratch under the same protocol as the original BioMod.

Module shuffling reduced the overall zero-shot integration score in all four evaluation settings (Supplementary Table S9). Averaging the displayed scores across the four settings, the mean overall score decreased from approximately 0.740 to 0.655. The mean biological-conservation score decreased by approximately 0.112, compared with a decrease of approximately 0.044 in the mean batch-mixing score. The largest reduction was observed for hPNC, where the overall score decreased from 0.665 to 0.486.

Cell-type annotation was also affected. Accuracy decreased on six of seven datasets, with mean accuracy decreasing from 0.864 to 0.827, while macro-F1 decreased on all seven datasets, with its mean decreasing from 0.812 to 0.726 (Supplementary Table S10). The largest accuracy drop occurred on MS (0.793 → 0.636). On Myeloid, accuracy increased from 0.615 to 0.627, while macro-F1 decreased from 0.645 to 0.339, indicating that the two metrics captured different aspects of annotation performance.

Shuffling reduced performance in both drug-response tasks: single-drug Pearson correlation decreased from 0.931 to 0.918, and drug-combination AUROC decreased from 0.681 to 0.628 (Supplementary Table S11). Its effect on perturbation prediction depended on the setting and metric. MSE decreased in Seen 0/2 (0.175 → 0.160) and Seen 0/1 (0.171 → 0.162), but increased in Seen 1/2 (0.180 → 0.189) and Seen 2/2 (0.122 → 0.149). Pearson correlation decreased in Seen 0/2 (0.871 → 0.857) and Seen 1/2 (0.876 → 0.852), but increased slightly in Seen 2/2 (0.953 → 0.954) and Seen 0/1 (0.876 → 0.878).

#### Post-encoder module-family ablation

To assess the relative contribution of each knowledge source, we performed a post-encoder ablation on the Zheng68K cell-type classification task. Unlike the preceding shuffled-module experiment, this analysis did not involve model retraining: all modules were first encoded with fully visible inputs, after which tokens from removed families, or tokens outside the retained families, were set to zero before task-specific pooling and classification. Input gene expression was unchanged, including genes shared between families; Reactome-only retained the 491 encoded Reactome tokens.

The complete-model accuracy was 0.839 in the post-encoder evaluation pipeline (Supplementary Table S12). Among the reported family-masking configurations, removing Reactome caused the largest accuracy decrease: accuracy fell to 0.764 after Reactome masking, while the configuration labeled Reactome-only had the highest single-family accuracy (0.713). By comparison, removing Hallmark, Hotspot or WGCNA modules resulted in accuracies of 0.802, 0.819 and 0.821, respectively. Pairwise retention confirmed the same trend: all Reactome-containing pairs outperformed the corresponding pairs without Reactome, with Reactome combined with Hotspot achieving the highest two-family accuracy (0.788). The complete four-family representation nevertheless outperformed every single-family and two-family configuration, so the unmasked model performed best among the tested configurations. Reactome accounts for 491 of the 660 modules; these interventions do not establish a size-matched comparison of knowledge sources.

In summary, the effect of biological gene-to-module assignments depends on the task and metric. When module count, module-size distribution, architecture and training protocol were held fixed, shuffling reduced zero-shot integration scores, macro-F1 on all seven annotation datasets, accuracy on six of seven annotation datasets, and performance on both drug-response tasks. Perturbation-prediction results were mixed across settings and metrics. Because both models used the same module-token architecture, this control tests biological organization within that architecture; it does not compare module tokens with gene tokens or test whether module tokenization reduces the required model size. In the post-encoder source-family analysis on Zheng68K, retaining Reactome alone gave the highest single-family accuracy and removing it caused the largest accuracy decrease, while the complete four-family representation achieved the highest accuracy.

### 2.8 Biological interpretation of learned module representations

Because modules are named, we could examine which gene programs contribute to cell-type classification and how module representations vary across datasets. We used pre-encoder module-importance rankings for canonical marker validation and post-encoder progressive masking to assess classifier dependence on contextualized module tokens. We also performed representational similarity analysis (RSA) with controls for module composition. These analyses separate whether classifier decisions can be traced to known gene programs from whether module embeddings align with pathway-specific semantics.

#### Module importance aligns with known cell-type marker programs

We next examined the modules used by a Zheng68K cell-type classifier [23]. For each of 13,690 test cells, we set the input expression values of each module’s genes to zero, reran the encoder and classifier, and measured the decrease in the correct-class logit. Importance scores were averaged within each cell type. Of 61 canonical cell-type–marker entries, 59 mapped to at least one module; the other two were excluded from the rank summaries. Among the 59 mapped entries, 41 (69.5%) had a containing module in the top 5% for the corresponding cell type, with a median best-module rank percentile of 0.91% (Fig. 6b; Supplementary Fig. S2d and Table S13). Because markers can occur in several modules, we compared this count with a membership-preserving random reference: randomly selecting 33 modules per cell type covered 15.1 entries on average (25.7%), and none of 20,000 random draws reached the observed count (one-sided empirical *P* ≈ 5.0×10^−5^). The pre-encoder top-10% count was 44/59 (74.6%). LYZ, S100A8 and S100A9 in monocytes, and CD8A, CD8B and GZMB in cytotoxic T cells, each had a best containing-module rank of 1. These are module ranks, not individual-gene importance ranks. This correspondence supports the biological readability of the module rankings, although marker membership in an overlapping gene set does not establish the causal contribution of an individual gene or activation of the entire pathway.

**Figure 6.**
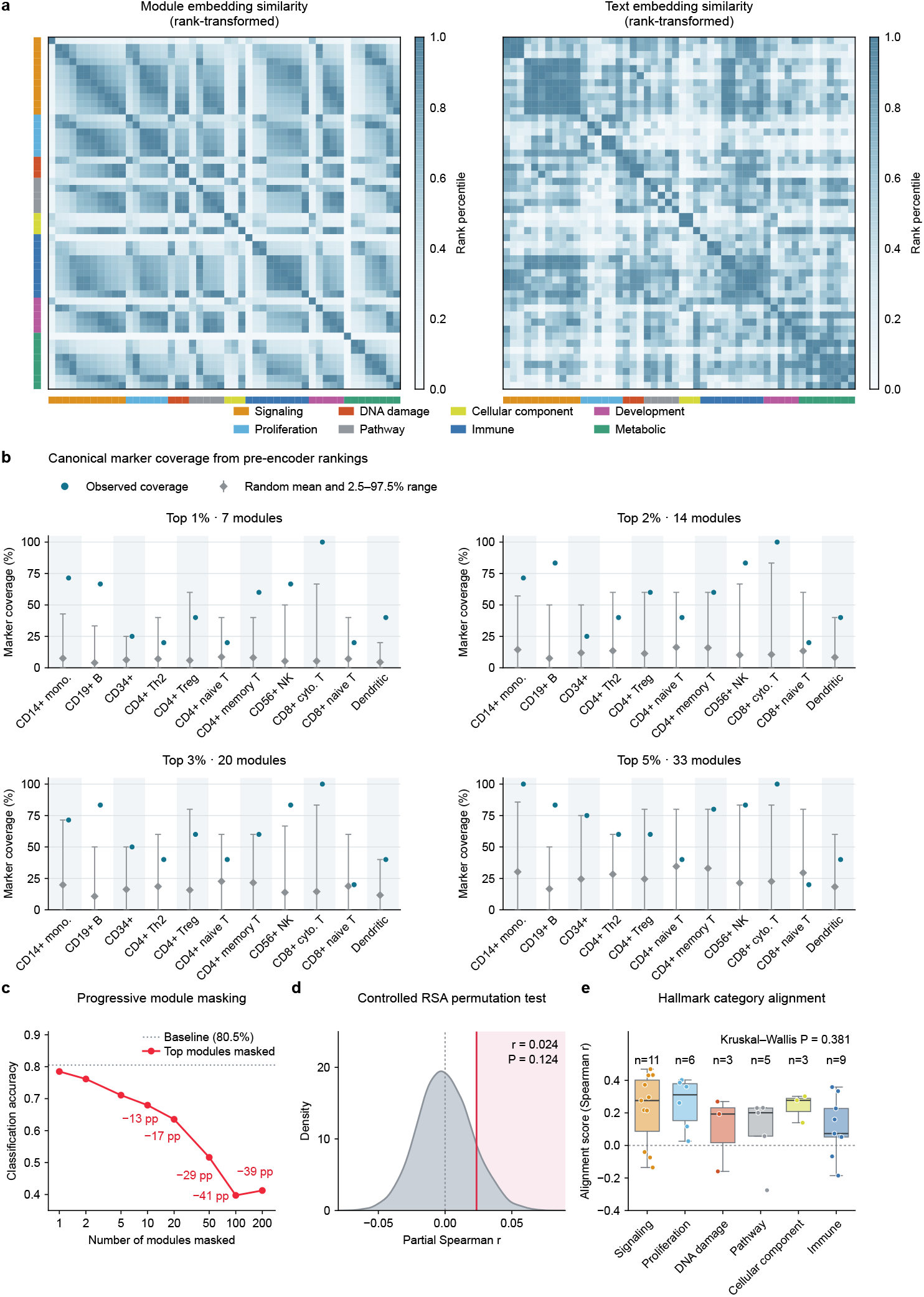
Module-level analyses of BioMod representations. **(a)** Comparison of pairwise similarity structures between module embeddings and pathway text embeddings across Hallmark [10] pathways. Both similarity matrices are rank-transformed; colored annotation bars indicate biological process categories. **(b)** Fractions of mapped canonical cell-type–marker entries covered by the top-ranked modules for each cell type, using pre-encoder importance rankings. An entry is covered when at least one module containing its marker is selected. The four panels show the top 1%, 2%, 3% and 5% of modules, corresponding to 7, 14, 20 and 33 of 660 modules, respectively. Blue circles indicate observed coverage. Gray diamonds and vertical error bars indicate the mean and 2.5th–97.5th percentiles of coverage from 20,000 random selections of the same number of modules. Random selections preserve the original module–gene memberships, accounting for markers that belong to multiple modules. **(c)** Classification accuracy under progressive post-encoder masking of top-ranked modules. Modules were ranked by post-encoder importance averaged across the 11 cell types. Selected module tokens were zeroed after a fully visible encoder pass. The dashed horizontal line indicates the unmasked baseline; masking 200 modules yielded an accuracy of 0.413. **(d)** Controlled module–text representational similarity analysis (RSA) for the pretraining subset, adjusted for module-source family, effective module size and gene-set Jaccard overlap. The gray distribution shows the permutation reference obtained by permuting text assignments within source families. The red vertical line indicates the observed partial Spearman correlation, *r* = 0.0236 (*P* = 0.124). **(e)** Distributions of per-Hallmark module–text alignment scores across the six displayed biological process categories. Points represent individual Hallmark pathways, and *n* denotes the number of pathways in each category. The between-category difference was not significant (Kruskal–Wallis *P* = 0.381).

Progressive masking examined dependence on contextualized module tokens. Modules were ranked by post-encoder importance averaged across the 11 cell types, and selected encoded tokens were set to zero after a fully visible encoder pass. In the same experiment, unmasked accuracy was 0.839 and decreased to 0.413 after masking the top 200 module tokens (Fig. 6c). Masking the bottom 200 retained 0.793 accuracy, while random masking retained 0.763 on average across five trials. No input genes were deleted or zeroed in any of these conditions. The larger drop for top-ranked tokens indicates greater classifier dependence on those representations; it does not measure input gene removal or biological pathway inhibition.

#### Module geometry is stable across datasets, but controlled text alignment is weak

We examined the same 541 pathway-based modules, comprising 50 Hallmark gene sets [11] and 491 Reactome pathways [12], in 50,000 pretraining cells and 10,000 cells from the independent pancreas dataset [25]. Module representational dissimilarity matrices were concordant across the two datasets (Spearman *r* = 0.900), including after adjustment for module-source family, effective module size and gene-set Jaccard overlap (*r* = 0.851). Across the full 660-module vocabulary, source-family organization was also present in the separately pretrained shuffled-module control and the initialized model: 10-nearest-neighbor family purity on pancreas was 0.755 for the main model, 0.740 for the shuffled control and 0.712 for the initialized model. The cross-dataset stability therefore reflects reproducible module geometry and does not by itself indicate a training-specific biological advantage.

The unadjusted module–text RSA [37] was positive in both the pretraining subset (*r* = 0.366) and pancreas (*r* = 0.407). However, after controlling for source family, effective module size and gene-set overlap, residual alignment was small and not significant: partial *r* = 0.0236, *P* = 0.124 in the pretraining subset (Fig. 6d), and partial *r* = 0.0264, *P* = 0.120 in pancreas, using 5,000 family-preserving permutations. The similarity heatmaps therefore describe representation structure rather than independently validating pathway semantics (Fig. 6a). Alignment scores also did not differ significantly among the six process categories displayed in Fig. 6e (Kruskal–Wallis *H* = 5.295, *P* = 0.381). After these controls, the raw correlations are largely explained by module composition, and we found no evidence of pathway-level semantic alignment beyond the tested factors.

## 3 Discussion

BioMod shows that biological module tokens can support cell-type annotation, zero-shot integration, perturbation-response prediction and drug-response prediction with approximately 50 million parameters. Its performance relative to scLong and Cell2Sentence shows that a compact module model is feasible, but does not demonstrate that tokenization reduces the required parameter count. Prior work has used pathway tokens for supervised annotation in TOSICA [14] and examined tokenizer design in HEIMDALL [16]. BioMod combines expression-conditioned tokens from curated pathways and data-derived programs with masked reconstruction at gene resolution, and we evaluated its transfer across four task types.

The shuffled-module comparison provides evidence about gene grouping within this architecture. Biological assignments improved integration, annotation macro-F1 and drug-response prediction under the tested protocols. Their effects on perturbation prediction were mixed, despite the strong performance of the complete model against baseline pipelines. Module representations can therefore support gene-level prediction, but biological grouping does not provide a consistent advantage for every setting or metric. These controls compare two pretrained module models; they do not isolate the contribution of pretraining itself or compare module and gene tokens under a matched training budget.

The correspondence with canonical markers supports the biological relevance of the input gene-set rankings. Pre-encoder masking measures sensitivity to the selected gene sets, including their effects on overlapping modules. Differences in the number of unique genes removed can also influence top-ranked, bottom-ranked and random comparisons. The family-masking results describe sensitivity under the specified interventions; the larger effect of Reactome is not a size-matched measure of source quality. Module geometry was reproducible across datasets, while controlled module–text alignment was small and nonsignificant. These analyses support a correspondence with known gene programs, but not causal attribution to individual pathways.

Module tokenization moves gene-level aggregation before global encoding. The encoder processes 660 tokens, or 165 visible tokens during pretraining, while reconstruction supervision remains at gene resolution within masked modules. The vocabulary covers 12,895 of 27,874 input genes, and genes outside the modules do not enter token construction. Module overlap also means that masked reconstruction targets may still be represented in visible modules. TSSA [20] and the shorter encoder sequence reduce architectural costs. Direct measurement on one GPU showed higher forward throughput, lower batch-size-1 latency and no out-of-memory failure up to batch size 256 for BioMod than for the compared models (Fig. 1e), although these timings cover only the representation forward pass and do not include preprocessing, data transfer or training cost. BioMod is smaller than scLong and Cell2Sentence, close in parameter count to scGPT, and larger than Geneformer and scBERT in Supplementary Table S4.

Further comparisons should match training data, parameter count, optimization budget and downstream heads to distinguish the contributions of tokenization, architecture and pretraining. Gene-coverage analyses and controls matched for the amount of input removed would clarify the module-sensitivity results. The present benchmarks report performance under the stated splits; transfer to unseen donors, drugs and cell lines requires separate evaluation. Overall, biological module tokens are useful units for compact single-cell models, with benefits that depend on the task and evaluation setting.

## 4 Methods

### 4.1 Pretraining corpus

To increase the scale and tissue diversity available for self-supervised pretraining while limiting over-representation of highly abundant tissues, we constructed a new human single-cell corpus from the public single-cell database scBaseCount [21]. After restricting the release to human scRNA-seq profiles, the source release comprises 35,259 public human single-cell RNA-sequencing samples and 289,251,287 cells. We adopted the ordered 27,874-gene vocabulary released with scLong [7] to provide a stable Ensembl-based gene axis and to maintain compatibility with the pretrained Gene2Vec resources used by BioMod. The scLong vocabulary was constructed by first removing non-human genes and harmonizing identifiers to Ensembl IDs, and then taking the union of the 20,000 genes most frequently detected across its source corpus, 19,748 annotated protein-coding genes, and genes represented in Gene Ontology or Gene2Vec, followed by deduplication. This design retains broad protein-coding coverage and genes supported by functional or semantic priors, rather than relying on an unfiltered union of source-specific features.

To curate tissue-resolved data, 3,964 source tissue labels were assigned to organ/tissue categories using an audited lookup table. Samples with unmapped, unknown, ambiguous, or explicitly excluded labels were omitted. Cells were retained when `gene_count_Unique` ≥ 500 and `umi_count_Unique` ≥ 1,000, and samples were retained when at least 50 cells passed both filters. To reduce tissue imbalance, we applied a target cap of 10 million cells per organ at the whole-sample level. The resulting corpus contains 72,993,441 cells from 13,527 samples across 25 organ/tissue categories.

Each retained sample was aligned to the ordered scLong vocabulary. For a raw UMI count *x*_*g*_, preprocessing computed *y*_*g*_ = log(1 + *x*_*g*_*/*10,000). Within each nonzero cell, expression values were rescaled to a maximum of 10 when max_*g*_ *y*_*g*_ ≤ 10; values above 10 were otherwise clipped. The resulting sparse expression matrices were retained together with sample, tissue, barcode, and global cell-index annotations. No samples were shared between the pretraining data, the module-discovery atlas and the evaluation datasets.

### 4.2 Module vocabulary construction

BioMod represents the transcriptome using a fixed, overlapping vocabulary of 660 biological modules assembled from four complementary sources: 50 MSigDB Hallmark modules, 491 Reactome modules, 73 Hotspot modules, and 46 postprocessed hdWGCNA modules. Modules are allowed to overlap within and across sources by design.

Curated pathway modules were obtained from MSigDB Hallmark version 2025.1 [10, 11] and Reactome 2022 [12]. All 50 Hallmark gene sets were included, comprising 7,322 source-level gene memberships and 4,384 unique gene symbols. Of the 1,816 Reactome pathways initially considered, 491 passed the predefined size filter of 50–500 genes, comprising 55,954 gene memberships and 9,745 unique gene symbols.

To construct data-driven modules, we assembled a compact, cancer-diverse atlas that enabled computationally tractable module discovery. Human scRNA-seq datasets spanning a broad range of cancer types and associated tissues were collected from CELLxGENE [1]. To reduce the dominance of highly represented cancer-associated tissues while preserving broad disease coverage, cells were downsampled within each harmonized tissue category to a maximum of 10,000 cells, followed by standard cell- and gene-level quality control. The resulting atlas contained 262,438 cells and 33,358 genes , spanning 31 tissue groups. Major populations included epithelial (68,511 cells), T cells (61,195), myeloid cells (31,710), malignant cells (31,593), fibroblasts (15,807), stem/progenitor cells (12,210), endothelial cells (7,647), and B cells (6,984).

For Hotspot [17], an integer-valued approximate count matrix was reconstructed by reversing the log transformation, estimating a cell-specific scaling factor from nonzero expression values, and rounding the rescaled values. Hotspot was fitted with the depth-adjusted negative-binomial noise model using per-cell total counts and Harmony embedding [28]. An unweighted *k*-nearest-neighbor graph was constructed with *k* = 30. Autocorrelation statistics were computed for all genes. Genes with nominal *P <* 0.05 were ranked by autocorrelation *Z*-score and limited to at most 5,000 candidates; 3,427 genes passed this criterion. Pairwise local-correlation *Z*-scores were then calculated, undefined entries were set to zero, and the matrix was symmetrized before hierarchical clustering. Modules were called with a minimum size of 15 genes, and an FDR threshold of 0.05, yielding 73 modules containing 2,489 genes (15–123 genes per module).

For hdWGCNA [18], the same 262,438 cell atlas was converted to a Seurat object. Genes detected in at least 5% of cells were selected, and metacells were constructed within cell-type-by-dataset groups in the Harmony space using *k* = 25, `max_shared=10`, and a minimum group size of 100 cells. A signed co-expression network was constructed from normalized metacell expression using `deepSplit=4`, `minModuleSize=20`, and `mergeCutHeight=0.05`. After removing genes with identifiers beginning with `ENSG`, the final postprocessed collection comprised 46 hdWGCNA-derived modules containing 3,850 genes (10–497 genes per module).

The four source collections (50 Hallmark, 491 Reactome, 73 Hotspot and 46 hdWGCNA modules) were merged into a single vocabulary of 660 modules. Because the sources use different gene nomenclatures, we mapped all identifiers to the Ensembl-based gene vocabulary adopted by the pretraining corpus [7]. Gene symbols are first converted to Ensembl IDs via an HGNC reference table and then indexed into the model vocabulary. Genes without a valid mapping were removed. After mapping, all 660 modules remained nonempty, comprising 68,985 gene memberships and 12,895 unique Ensembl genes. Modules were allowed to overlap by design, enabling the same gene to be represented in multiple functional contexts.

The module collection covers approximately 46.3% of the 27,874-gene input vocabulary. Genes outside the collection do not directly contribute to module-token construction or masked reconstruction. The module definitions and their ordering were fixed throughout pretraining and downstream evaluation.

### 4.3 Model architecture and pretraining

BioMod applies two levels of computation. Within each module, a local step combines gene embeddings with expression values to construct one module token. The global encoder then models relationships only among the resulting 660 module tokens. This shortens the sequence used for global modeling; it does not eliminate gene-level computation before module construction.

#### Combining genes within modules

Within each module, the normalized expression value of each gene is multiplied by its pretrained Gene2Vec embedding. For any gene *g* in module *p*_*i*_, the expression value *x*_*g*_ and embedding *e*_*g*_ (initialized from Gene2Vec [19], dimension 200) are combined as:

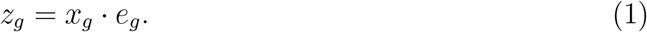

The Gene2Vec embedding is initialized from pretrained gene co-expression patterns and fine-tuned during training. The resulting feature *z*_*g*_ is mapped to the module-token dimension through a shared nonlinear projection, *h*_*g*_ = *W*_2_*σ*(*W*_1_*z*_*g*_ + *b*_1_) + *b*_2_.

Each projected gene feature receives a learned score. Let *s*_*g*_ = **w**^⊤^*h*_*g*_; the normalized weight *α*_*g*_ and module token **t**_*i*_ are defined as:

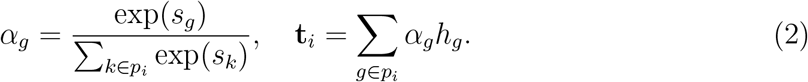

This learned weighting allows different genes to contribute by different amounts to the module representation of each cell. The resulting sequence of *L* module tokens, **X**_0_ = [**t**_1_ + *E*_*pos*,1_, … , **t**_*L*_ + *E*_*pos,L*_], is passed to the encoder, where *E*_*pos*_ denotes module identity embeddings.

#### Encoder

The encoder uses token-statistical self-attention (TSSA) [20] to model cross-module dependencies with linear complexity. TSSA replaces the explicit *O*(*L*^2^) attention matrix with prefix-state aggregation over a single linear projection. For input sequence **X** ∈ ℝ^*L*×*D*^, a shared projection produces per-head states 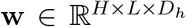. A cumulative normalisation term and a bias-corrected relevance score are computed as:

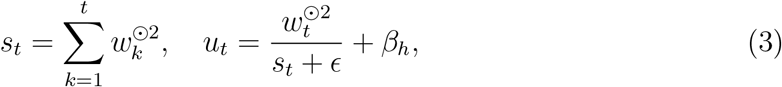

where ⊙2 denotes element-wise square, *ϵ* is a small constant for numerical stability, and *β*_*h*_ is a learnable per-head bias. Head-wise gating weights are obtained through a temperature-scaled softmax across heads:

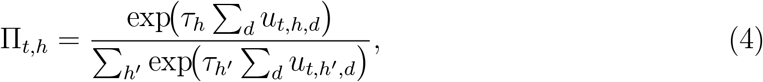

where *τ*_*h*_ is a learnable temperature parameter per head. The output is formed using a cumulative gated state *d*_*t*_ and a suppression factor *a*_*t*_:

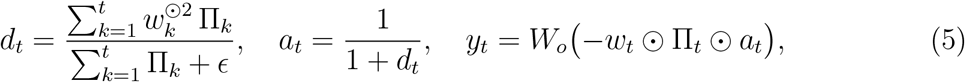

where *W*_*o*_ is a learned output projection. The entire computation involves only cumulative sums and element-wise operations, yielding *O*(*L*) time and memory complexity. The complete model used for the reported experiments contains approximately 50 million trainable parameters.

#### Masking and decoder

We adopt a drop-mask strategy inspired by masked autoencoders (MAE) [38], in which masked module tokens are removed from the encoder input rather than replaced by mask tokens. For each input cell, i.i.d. uniform noise *n*_*i*_ ∼ *U*(0, 1) is sampled for each module token, and all *L* = 660 tokens are ranked accordingly. Following MAE, a high masking ratio of *r* = 0.75 retains only 25% of the module tokens (*L*_*keep*_ = 165) as visible inputs. Let the masking indicator *m*_*i*_ ∈ {0, 1}, where *m*_*i*_ = 1 denotes masked and *m*_*i*_ = 0 denotes visible. The encoder output is:

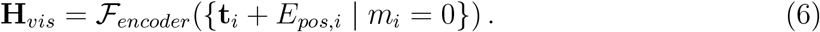

The encoder receives a small subset of module tokens. Because modules overlap, genes in a masked module may also occur in visible modules; the module masking ratio therefore does not equal the fraction of gene-expression information withheld. After masking, the visible tokens are randomly shuffled, so the encoder learns relationships among tokens without relying on a fixed module order.

The decoder reconstructs masked-module expression vectors from encoded visible states using pure-memory cross-attention, in which visible token states serve as memory **M** and masked-module query embeddings serve as queries **Q**. Encoded visible states are passed to the decoder without additional positional embeddings or dropout. The decoder layer performs:

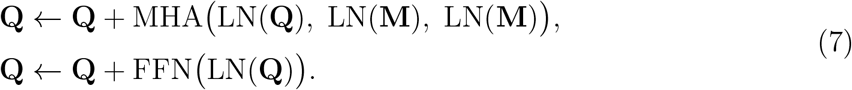

The decoder conditions masked-module reconstruction on the encoded visible states. Shared genes may remain represented in visible modules, so reconstruction can use both shared gene information and relationships across modules.

#### Reconstruction loss

The reconstruction loss is computed at gene resolution. For each masked module *p*_*i*_, the decoder output *h*_*i*_ is projected by a module-specific linear head:

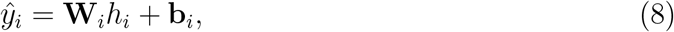

where 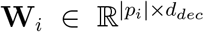 is a module-specific projection matrix whose output dimension matches the gene cardinality of module *p*_*i*_, so that reconstruction is constrained to the predefined module boundaries while preserving gene-level supervision.

For the *i*-th masked module *p*_*i*_ containing |*p*_*i*_| genes, the within-module mean squared error is:

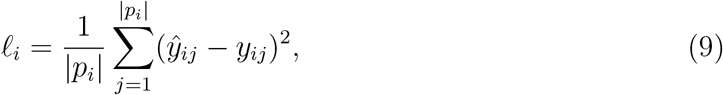

where *y*_*ij*_ and *ŷ*_*ij*_ are ground-truth and reconstructed expression values, respectively. The global reconstruction loss *ℒ*_*rec*_ is averaged across all masked modules:

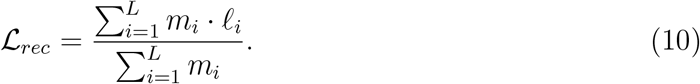

Because pathway gene cardinality varies widely (from tens to hundreds of genes), a naïve global gene-level MSE would weight gradient contributions in proportion to module size, biasing learning toward large pathways. The two-stage averaging in *ℒ*_*rec*_ normalizes first within each module and then across modules, assigning equal weight to each masked module’s mean reconstruction loss rather than weighting modules in proportion to their gene counts. This does not imply equal gradient magnitudes across modules.

#### Training details

Pretraining is distributed across 64 H100 GPUs with a per-device batch size of 75 and two-step gradient accumulation, yielding an effective global batch size of 9,600. Optimization uses Adam with an initial learning rate of 1×10^−4^, cosine annealing with warmup restarts, and mixed-precision training.

### 4.4 Downstream task adaptation

All downstream tasks share a common input pipeline. For an input expression profile 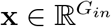, we first reorder it into the pretrained gene space *G* through Ensembl-ID mapping, yielding **x**^(*model*)^:

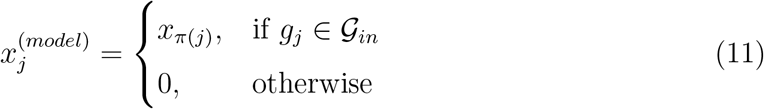

where *π*(*j*) denotes the index mapping of the original gene in the input vector. The reordered vector is passed through the BioMod backbone in fully visible mode (mask ratio 0), producing a module-level representation sequence **H** = [**h**_1_, … , **h**_*L*_] ∈ ℝ^*L*×*d*^. Depending on the task, the backbone is either fine-tuned end-to-end (cell-type annotation) or kept frozen as a feature extractor (drug-response prediction and zero-shot integration). For perturbation prediction, the backbone is frozen and module representations are projected back to gene-level features (Section 4.7).

### 4.5 Cell-type annotation

We evaluated cell-type annotation on seven benchmarks spanning diverse tissues and annotation granularities: Zheng68K [23] (68,450 PBMCs spanning 11 immune subtypes), lung, immune, MS, liver [22], hPancreas [14], and Myeloid [24]. We used two data-partitioning strategies depending on the original organization of each benchmark. For datasets provided as a single expression matrix without a predefined test set, cells were divided by stratified sampling into training, validation, and test sets at a ratio of 70%:10%:20%, preserving cell-type proportions across partitions. For datasets with predefined training and test partitions, the original test set was retained unchanged for final evaluation, whereas the provided training set was further divided by stratified sampling into training and validation subsets at a ratio of 90%:10%. No cells from the predefined test set were used for model selection or hyperparameter tuning.

The backbone was fine-tuned end-to-end. Module representations were combined into a single cell embedding, **h**_cell_, using learned weighted averaging: a two-layer scoring network assigns a relevance score to each module token, the scores are normalized with a softmax, and the tokens are summed using these weights. The resulting embedding was passed through dropout (*P* = 0.2) and then a three-layer classifier that produced the predicted class-probability distribution. The model was optimized using the cross-entropy loss:

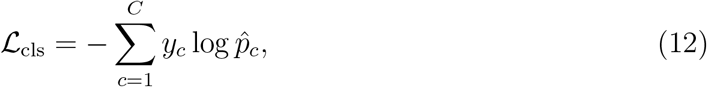

where *C* denotes the number of cell types, *y*_*c*_ is the one-hot-encoded ground-truth label, and 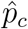 is the predicted probability for class *c*. The backbone and classification head were initialized with a learning rate of 1 × 10^−4^ and jointly optimized using AdamW with a weight decay of 1×10^−2^. A cosine-annealing learning-rate schedule was applied throughout fine-tuning.

### 4.6 Zero-shot batch integration

We used three benchmark atlases with substantial batch heterogeneity: the hPNC dataset [25] (16,382 cells, 19,093 genes, 6 sequencing batches, 14 cell subtypes), the BMMC dataset [25] (90,261 cells, 12 batches), and a human lung atlas [26] (32,493 cells, 14 patient-sample batches, 10 cell lineages). For the lung atlas, metrics are reported both with malignant cells retained and after malignant-cell removal. We retained all gene features without highly-variable-gene selection or additional cell filtering.

All backbone parameters were frozen during inference. Module-level representations **H** are reduced to a single cell vector **v** via max pooling:

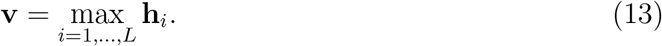

We evaluated embedding quality using the scIB metric suite [25]. A cell-cell *K*-nearest-neighbor graph is constructed from **v**, and two composite scores are computed. The Biological Conservation Score (avg_bio_) combines normalized mutual information (NMI), adjusted Rand index (ARI), and label-based average silhouette width (ASW) to quantify preservation of cell-subtype boundaries:

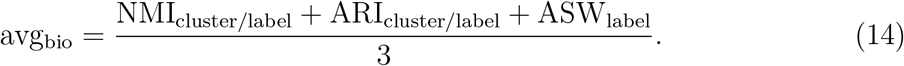

The Batch Mixing Score (avg_batch_) combines graph connectivity and batch-aware silhouette width to measure cross-batch mixing of cells sharing the same type:

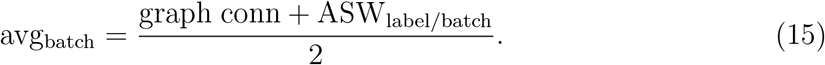

The overall integration score is a weighted average of the two components:

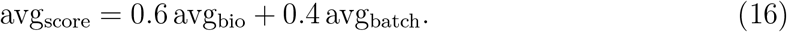

The resulting embeddings are visualized with UMAP [39] and compared against specialized batch-correction methods.

### 4.7 Genetic perturbation prediction

We integrated BioMod into the GEARS [30] framework as a pretrained single-cell encoder and evaluated it on the Norman dataset [29], which contains 91,205 cells, 5,045 genes, and 236 perturbation conditions (105 single-gene and 131 double-gene perturbations). Following the GEARS simulation split, we assigned 58,134, 6,792 and 26,279 cells to the training, validation and test sets, respectively. The test set was further divided into four generalization categories: unseen single perturbations (Seen 0/1) and combinatorial perturbations with varying degrees of novelty (Seen 0/2, Seen 1/2, and Seen 2/2).

#### Module-to-gene recovery

Because GEARS operates at gene resolution, BioMod module representations must be projected back to individual genes. Let the control-state expression vector be **x** ∈ ℝ^5045^ over the dataset’s gene set. BioMod first reorders the input into the pretrained gene vocabulary G, then outputs module representations **H** = [**h**_1_, … , **h**_*L*_] ∈ ℝ^*L*×*d*^ under fully visible mode. Let *S*(*k*) denote the set of modules associated with gene *k*; the recovered gene representation is

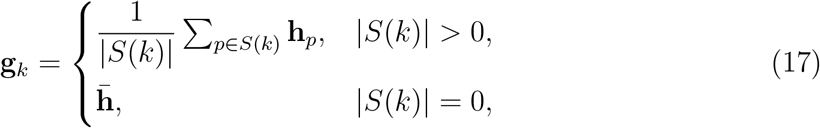

where 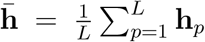 is the global module-mean representation. The recovered gene representations are then linearly projected to the GEARS hidden dimension.

#### GEARS perturbation encoder and decoder

Perturbation conditions are encoded as a gene-level binary indicator vector 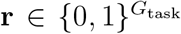, where *r*_*k*_ = 1 indicates that gene *k* is perturbed and *G*_task_ = 5,045 is the task gene set size. Following GEARS [30], each gene is assigned a learnable perturbation embedding that is refined by one layer of message passing over a GO similarity graph. Embeddings of all perturbed genes are aggregated into a condition-level representation **c**_pert_ = *f*_pert_(Σ_*k*_ *r*_*k*_**u**_*k*_), which is broadcast and added to all gene representations. A perturbation-position embedding based on *r*_*k*_ further distinguishes perturbed from unperturbed genes. The fused gene features are batch-normalized and passed to the GEARS decoder, which produces a gene-specific intermediate response via an MLP, aggregates all gene responses through a cross-gene MLP to capture inter-gene dependencies, and outputs the final prediction in residual form:

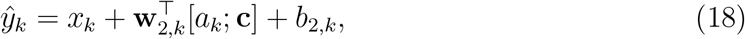

where *a*_*k*_ is the gene-specific intermediate scalar, **c** is the shared cross-gene state, and **w**_2,*k*_, *b*_2,*k*_ are gene-specific parameters. The model thus learns perturbation-induced expression change rather than absolute expression directly.

#### Loss and evaluation

Following GEARS [30], the training loss combines a quartic error term *ℒ*_pred_ and a direction-consistency term *ℒ*_dir_ that penalizes sign mismatches between predicted and observed expression changes relative to control:

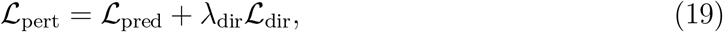

with *λ*_dir_ = 0.1. Both terms average first over genes within each perturbation condition and then across conditions, excluding genes that remain zero under the current perturbation. Training uses Adam with learning rate 1 × 10^−3^ and weight decay 5 × 10^−4^.

Evaluation metrics include global MSE and Pearson correlation, as well as MSE and Pearson correlation computed on the top 20 differentially expressed (DE) genes. For DE gene set *D*_*c*_ under condition *c*, the top-20 DE error is defined as

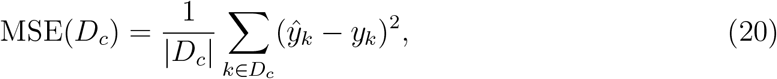

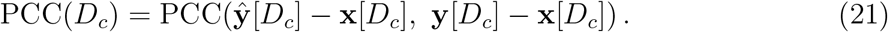

In addition to global metrics, we report subgroup-specific MSE and Pearson correlation under the simulation split for each of the four generalization settings (Seen 0/2, Seen 1/2, Seen 2/2, and Seen 0/1).

### 4.8 Drug-response prediction

We constructed a dual-branch cross-modal regression model that combines frozen BioMod cell-state representations with molecular graph features extracted by a graph neural network (GNN). Experiments used a joint CCLE [31] and GDSC [35] dataset. The benchmark initially matched 107,446 records and retained 94,314 after cancer-type filtering. Under the record-level split, approximately 66,020 records were assigned to training, 9,431 to validation and 18,863 to testing. Model parameters were fitted using the training set; the validation set was used for early stopping and checkpoint selection, and the test set was reserved for final evaluation. Splitting was by response record rather than by unique drug–cell-line pair; the training and test sets shared 222 cell lines, 223 drugs and 530 identical drug–cell-line pairs. The prediction target is the natural logarithm of half-maximal inhibitory concentration, ln(IC_50_). This evaluation concerns held-out response records; it does not directly test response in patients or generalization to unseen drug–cell-line pairs, drugs or cell lines.

#### Transcriptomic branch

BioMod serves as a frozen feature extractor. The 697 input genes were processed in the same way as for the other compared models. Of the 697 input genes, 689 (98.9%) mapped to the pretrained vocabulary and 605 (86.8%) occurred in the module collection. Unmatched columns were excluded and missing vocabulary positions were filled with zero. For each cell-line expression vector **x** ∈ ℝ^697^, expression values are padded and reordered via Ensembl ID mapping into the pretrained gene vocabulary G. The reordered vector is passed through the frozen backbone, and a global cell embedding is obtained by max pooling:

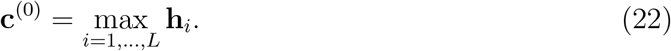

The global embedding is then transformed by a two-layer MLP with batch normalization and ReLU activations, yielding the final cell-state representation **c** ∈ ℝ^128^.

#### Drug-graph branch

Each small molecule is represented as an undirected molecular graph encoded by a three-layer graph convolutional network (GCN) [40] with hidden dimensions 256 → 256 → 128 and a residual connection in the second layer. The drug representation **d** ∈ ℝ^128^ is obtained by summing element-wise max pooling and mean pooling over the final-layer node features.

#### Cross-modal fusion and training

The cell representation **c** and drug representation **d** are concatenated into a joint vector **h** = [**d**; **c**] ∈ ℝ^256^, which is modulated by a sigmoid attention gate and decoded by an MLP (512 → 256 → 128 → 1) to predict ln(IC_50_). The model is trained with Huber loss (*δ* = 1.0) using AdamW [41] (learning rate 1 × 10^−3^; weight decay 1 × 10^−5^) with cosine-annealing scheduling and warm restarts. Training runs for up to 300 epochs with batch size 32 and early stopping (patience 30 epochs). Pearson correlation on the validation set was used for early stopping and checkpoint selection; the test set was reserved for final performance evaluation.

#### Drug-combination synergy prediction

We constructed a separate model for drug-combination synergy prediction. The transcriptomic branch again uses frozen BioMod with max pooling, followed by *L*_2_ normalization and a three-layer MLP (512 → 256 → 128) to produce the cell-state vector **c** ∈ ℝ^128^. Each drug in the pair is encoded by a shared three-layer GCNConv branch (78 → 156 → 312) with global max pooling, followed by two fully connected layers (312 → 156 → 128) to yield drug representations **d**_1_, **d**_2_ ∈ ℝ^128^. The three representations are concatenated into [**d**_1_; **d**_2_; **c**] ∈ ℝ^384^ and decoded by a two-layer MLP (256 → 128 → 2) to predict a binary synergy label. The model is trained with cross-entropy loss. We evaluated performance using the area under the receiver operating characteristic curve (AUROC).

### 4.9 Module embedding–pathway text alignment

We quantified the correspondence between BioMod module embeddings and pathway-level semantic descriptions using representational similarity analysis (RSA) [37]. The analysis included 541 pathway-based modules, consisting of 50 MSigDB Hallmark gene sets [10] and 491 Reactome pathways [12]. For each pathway module, we constructed a text field from the pathway name and, when available, the Reactome summary description. Text fields were encoded using S-PubMedBert-MS-MARCO [42], a biomedical sentence encoder trained for sentence-level semantic similarity, yielding a 768-dimensional text embedding for each pathway.

Module embeddings were obtained by running the pretrained BioMod encoder in fully visible mode (mask ratio set to 0) and, for each module, averaging its encoded token representation across cells. The primary analysis used module embeddings computed from a 50,000-cell subset of the pretraining corpus. To evaluate whether the result was datasetspecific, we repeated the analysis using module embeddings computed from an independent pancreas scIB [25] dataset.

For RSA, we computed pairwise cosine distance matrices for the module embeddings and the text embeddings. The structural correspondence between the two spaces was measured as the Spearman rank correlation between the upper-triangular entries of the two distance matrices.

The controlled RSA reported in Results adjusted for module-source family, effective module size and gene-set Jaccard overlap. Significance was assessed using 5,000 permutations of text assignments within source families.

For Hallmark-level analysis, we restricted the comparison to the 50 MSigDB Hallmark modules [10]. Per-Hallmark alignment scores were computed by correlating each Hall-mark’s similarity profile to all other Hallmarks in module space with its corresponding similarity profile in text space. Hallmarks were further grouped into biological process categories, and category-level differences were tested using the Kruskal–Wallis test. To assess granularity effects, RSA was repeated at four levels: individual Reactome pathways [12], Reactome top-level categories, Reactome super-categories and MSigDB Hallmarks [10]. For category-level analyses, module embeddings and text embeddings were averaged within each pathway category before computing pairwise distances.

### 4.10 Module importance and marker validation

We assessed decision-level interpretability by measuring the contribution of each module to cell-type classification. The analysis used the BioMod model fine-tuned on the Zheng68K [23] cell-type annotation benchmark. The held-out test set contained 13,690 cells from 11 annotated immune cell types.

Progressive masking used a separate post-encoder analysis. Modules were ranked by post-encoder importance averaged across the 11 cell types. After one fully visible encoder pass, selected module tokens were set to zero before pooling and classification, with 1, 2, 5, 10, 20, 50, 100 or 200 tokens masked in top-ranked, bottom-ranked or random order. The input expression profile was unchanged. Random masking was averaged across five trials.

We used permutation controls to assess whether the observed importance structure exceeded random baselines. In the label-shuffle control, cell-type labels were randomly permuted and per-class importance summaries were recomputed 1,000 times from the stored per-cell perturbation scores. Empirical *P* values were computed as the fraction of null summaries greater than or equal to the observed statistic. We also performed a module-shuffle control in which module identities were permuted before recomputing importance; because this control requires repeated model evaluation, it was run for 10 permutations and was interpreted as a qualitative control rather than as a high-resolution *P* -value estimate.

Canonical marker validation mapped each cell-type–marker entry to all modules containing that gene. The best containing-module rank was recorded for the corresponding cell type. To evaluate top-5% coverage, 33 of the 660 modules were sampled uniformly without replacement for each cell type, retaining the original marker-to-module memberships. Across 20,000 random draws (seed 20260912), the mean coverage was 15.135 of the 59 mapped entries (25.7%). The pre-encoder ranking covered 41 entries; no random draw reached this count. The one-sided empirical probability was computed as (*b* + 1)*/*(*B* + 1), with *B* = 20,000 and *b* the number of random counts at least as large as the observed count. This reference accounts for overlapping marker memberships but does not match modules by size.

As an additional annotation comparison, genes contained in the top-ranked modules for each cell type were tested for enrichment against PanglaoDB [43] marker sets using one-sided hypergeometric tests with multiple-testing correction across database cell-type sets.

### 4.11 Baseline training and statistical analysis

For each downstream task, baseline foundation models with published task-specific pipelines (e.g., scGPT [3] and scFoundation [5] for perturbation prediction) were trained with their original code. Models without task-specific implementations were adapted using the same framework as BioMod: the GEARS-based pipeline for perturbation prediction, or the same dual-branch architecture (with the transcriptomic encoder replaced) for drug-response prediction. Task-specific baselines (GEARS [30], DeepCDR [34], DeepDDS [36]) were trained with their original implementations. Early stopping was applied when validation performance began to decrease. All models were evaluated using the same data splits and evaluation protocols. Cell-type annotation tables report means and standard deviations for BioMod and baselines where available. Drug-response tables report baseline means across five runs without standard deviations, whereas BioMod values are shown as mean ± s.d. Other results are reported as specified in the corresponding tables. All experiments, including those for the baselines, were repeated five times with different random seeds.

Detailed performance results for BioMod and each baseline across the downstream tasks are provided in the Supplementary Tables.

## Supporting information

supplemental for paper

## Supplementary information

Supplementary information is provided as a separate file, containing supplementary notes, benchmark summaries, dataset descriptions, and extended statistical tables supporting the analyses reported in the main manuscript.

## Acknowledgements

This work was supported by the Shenzhen Loop Area Institute under Grant No. FPF10120250014 and Contract No. SLAI2026020007, and by the Joint Laboratory for Single-Cell Datasets and Foundation Models, jointly established by the Shenzhen Research Institute of Big Data and Shenzhen Raymind Biotechnology Co., Ltd.

## Data availability

The pretraining corpus was curated from scBaseCount [21]. The separate atlas used to derive Hotspot and hdWGCNA modules was obtained from CELLxGENE [1]. Downstream evaluation datasets are publicly available and cited in the Methods section.

## Code availability

The pretrained model weights and inference code are available at https://huggingface.co/ctcatlas/BioMod.

