## supplemental for paper for "BioMod: Biological Module Tokenization for Single-Cell Foundation Modeling"

[4]Shenzhen Ruimai Biotechnology Co., Ltd.

### Supplementary Methods

This section provides additional implementation details, dataset-processing choices, model hyperparameters and evaluation protocols that support the analyses reported in the main manuscript.

#### S1.1 Dataset preprocessing

The pretraining corpus was assembled from public human single-cell RNA-sequencing profiles in scBaseCount [1]. The human subset of the source release comprised 35,259 samples and 289,251,287 cells. We adopted the ordered 27,874-gene vocabulary released with scLong [2]. This vocabulary was constructed by removing non-human genes, harmonizing identifiers to Ensembl IDs, and taking the union of the 20,000 genes most frequently detected across the scLong source corpus, 19,748 annotated protein-coding genes [3], and genes represented in Gene Ontology [4] or Gene2Vec [5], followed by deduplication.

To curate tissue-resolved data, 3,964 source tissue labels were assigned to organ/tissue categories using an audited lookup table. Samples with unmapped, unknown, ambiguous, or explicitly excluded tissue labels were omitted. Cells were retained when they had at least 500 detected genes and 1,000 UMIs, and samples were retained when at least 50 cells passed both filters. To reduce tissue imbalance, a target cap of 10 million cells per organ was applied at the whole-sample level. The resulting corpus contained 72,993,441 cells from 13,527 samples across 25 organ/tissue categories.

Each retained sample was aligned to the ordered scLong gene vocabulary. For a raw UMI count $x_g$, preprocessing computed $y_g = \log(1 + x_g/10{,}000)$. Let $m = \max_g y_g$ for a given cell. When $0 < m \leq 10$, expression values were rescaled as $10y_g/m$; when $m > 10$, values above 10 were clipped to 10. The resulting sparse expression matrices were retained together with sample, tissue, barcode, and global cell-index annotations. The final pretraining data scale was approximately 73.0 million cells, as reported in Supplementary Table S4.

#### S1.2 Model configuration

The pretrained BioMod backbone was implemented in the module-based scLong training framework [2] and launched with PyTorch distributed training. The training script used the scLong gene vocabulary, a biological module definition file containing 660 modules, Ensembl gene mapping [3] and pretrained Gene2Vec features [5]. The command-line configuration used for the final pretraining run is summarized below.

For pretraining, each worker loaded sharded expression matrices from the training data store, mapped genes to the fixed gene vocabulary, and constructed module tokens using the same module definitions across all cells. The model was optimized for 30 epochs with a high module masking ratio ($r = 0.75$) [7], gene-level reconstruction targets and the module-mean loss. The validation pass used a larger per-device batch size because no backward pass was required. Distributed training was coordinated with `torchrun`.

**Supplementary key hyperparameters for BioMod.** The table summarizes the main architectural and optimization hyperparameters used for final pretraining.

| Hyperparameter | Value |
|---|---|
| Input gene space | scLong gene vocabulary [2] |
| Module vocabulary | 660 biological modules |
| Gene–expression fusion | Element-wise multiplication of expression values and Gene2Vec embeddings [5] |
| Encoder backbone | 2-layer TSSA encoder [6] |
| Encoder hidden dimension | 1024 |
| Number of attention heads | 16 |
| Module mask ratio | 0.75 |
| Visible module ratio | 0.25 |
| Decoder type | Asymmetric cross-attention decoder |
| Decoder depth | 1 layer |
| Decoder embedding dimension | 256 |
| Reconstruction target | Gene-level reconstruction within masked modules |
| Pretraining loss | Module-mean reconstruction loss |
| Training epochs | 30 |
| Initial learning rate | $1 \times 10^{-4}$ |
| Per-device batch size | 75 |
| Gradient accumulation steps | 2 |
| Validation batch size | 175 |
| Validation interval | Every 30,000 training steps |

### Supplementary Results

This section provides extended quantitative results and additional analyses for BioMod across downstream tasks.

#### S2.1 Cell-type annotation

We evaluated BioMod on seven cell-type annotation benchmarks, including the Zheng68K, Kim Lung, MS, Liver, Immune, Myeloid and hPancreas datasets [8–12]. Across five independent runs, BioMod achieved mean accuracies of 0.839, 0.986, 0.793, 0.913, 0.948, 0.615 and 0.956 on these datasets, respectively, with corresponding mean macro-F1 scores of 0.752, 0.962, 0.679, 0.918, 0.930, 0.645 and 0.797 (Supplementary Table S1). Kim Lung had the highest accuracy and macro-F1 among the seven datasets. Accuracy and macro-F1 were also high on Liver and Immune. BioMod achieved high accuracy on hPancreas, although its macro-F1 was comparatively lower. Performance on MS and Myeloid was more modest. Annotation performance therefore varied across datasets despite using the same pretrained module-level backbone.

We next compared BioMod with representative single-cell foundation models [13–16] on datasets for which baseline results were available. The comparisons were grouped by dataset in Supplementary Table S2. On Zheng68K, BioMod achieved the highest accuracy and macro-F1 among the compared methods, reaching 0.839 and 0.752, respectively. Geneformer achieved an accuracy of 0.834 and a macro-F1 of 0.733. On Liver, BioMod

Table S1: **Cell-type annotation performance of BioMod across benchmark datasets.** Accuracy and macro-F1 are reported as mean ± s.d. across five independent runs. Macro-F1 denotes the unweighted mean of the per-class F1 scores.

| Dataset | Accuracy | Macro-F1 |
|---|---|---|
| Zheng68K | $0.839 \pm 0.005$ | $0.752 \pm 0.007$ |
| Kim Lung | $0.986 \pm 0.002$ | $0.962 \pm 0.005$ |
| MS | $0.793 \pm 0.014$ | $0.679 \pm 0.015$ |
| Liver | $0.913 \pm 0.011$ | $0.918 \pm 0.008$ |
| Immune | $0.948 \pm 0.003$ | $0.930 \pm 0.003$ |
| Myeloid | $0.615 \pm 0.015$ | $0.645 \pm 0.016$ |
| hPancreas | $0.956 \pm 0.004$ | $0.797 \pm 0.017$ |

achieved the highest macro-F1 of 0.918, while its accuracy of 0.913 was slightly below the joint highest value of 0.915 reported for scGPT and Geneformer. On hPancreas, BioMod achieved an accuracy of 0.956 and the highest macro-F1 of 0.797. scGPT obtained the highest accuracy of 0.973 and the second-highest macro-F1 of 0.757, followed by Geneformer with a macro-F1 of 0.752. On Myeloid, BioMod achieved the highest macro-F1 of 0.645 and an accuracy of 0.615. Its accuracy exceeded those of Cell2Sentence and scFoundation, but remained below those of scGPT and Geneformer. The relative ranking of BioMod therefore depended on both the dataset and the evaluation metric, with the clearest gains appearing in macro-F1.

The supplementary annotation results show a dataset-dependent pattern. BioMod performed best on Kim Lung and remained high on Liver and Immune, while hPancreas showed high overall accuracy with a lower macro-F1. MS and Myeloid were more difficult. On Myeloid, BioMod had the highest macro-F1 among the compared methods despite not attaining the highest accuracy, indicating that accuracy alone understated its class-balanced performance. Conversely, the gap between accuracy and macro-F1 on hPancreas shows that high overall accuracy can coincide with less even performance across cell types. Reporting both metrics is therefore necessary for these heterogeneous annotation benchmarks.

#### S2.2 Zero-shot integration

BioMod was evaluated in four zero-shot integration settings: Kim Lung with malignant cells retained, hPNC, BMMC and Kim Lung after malignant-cell removal [9, 17]. Biological conservation, batch mixing and the weighted overall score are reported in Supplementary Table S3 as mean and s.d. across five independent runs. In the Kim Lung setting, the biological-conservation score was $\text{avg}_{\text{bio}} = 0.647 \pm 0.010$, the batch-mixing score was $\text{avg}_{\text{batch}} = 0.982 \pm 0.000$, and the weighted overall score was $\text{avg}_{\text{score}} = 0.781 \pm 0.006$. After malignant-cell removal, the corresponding scores were $\text{avg}_{\text{bio}} = 0.761 \pm 0.020$, $\text{avg}_{\text{batch}} = 0.986 \pm 0.000$ and $\text{avg}_{\text{score}} = 0.851 \pm 0.012$. A compact comparison with Cell2Sentence [16] is included below the complete BioMod metric table. BioMod achieved the higher overall integration score on Kim Lung, hPNC and Kim Lung without malignant cells. On

Table S2: **Comparison of cell-type annotation performance across benchmark datasets.** Accuracy and macro-F1 are reported for BioMod and baseline models on benchmark datasets with available comparison results. Values are shown as mean ± s.d. where available, and BioMod values are means across five independent runs. The best value for each metric within each dataset is shown in bold.

| Dataset | Method | Accuracy | Macro-F1 |
|---|---|---|---|
| Zheng68K | scGPT | $0.794 \pm 0.008$ | $0.691 \pm 0.006$ |
| | scFoundation | $0.756 \pm 0.010$ | $0.605 \pm 0.002$ |
| | Geneformer | $0.834 \pm 0.002$ | $0.733 \pm 0.011$ |
| | Cell2Sentence | $0.650 \pm 0.012$ | $0.530 \pm 0.014$ |
| | BioMod | $\mathbf{0.839} \pm 0.005$ | $\mathbf{0.752} \pm 0.007$ |
| Liver | scGPT | $\mathbf{0.915} \pm 0.006$ | $0.831 \pm 0.005$ |
| | scFoundation | $0.626 \pm 0.099$ | $0.513 \pm 0.017$ |
| | Geneformer | $\mathbf{0.915} \pm 0.010$ | $0.770 \pm 0.007$ |
| | Cell2Sentence | $0.716 \pm 0.019$ | $0.688 \pm 0.020$ |
| | BioMod | $0.913 \pm 0.011$ | $\mathbf{0.918} \pm 0.008$ |
| hPancreas | scGPT | $\mathbf{0.973} \pm 0.007$ | $0.757 \pm 0.047$ |
| | scFoundation | $0.795 \pm 0.092$ | $0.476 \pm 0.078$ |
| | Geneformer | $0.970 \pm 0.010$ | $0.752 \pm 0.034$ |
| | Cell2Sentence | $0.963 \pm 0.003$ | $0.708 \pm 0.030$ |
| | BioMod | $0.956 \pm 0.004$ | $\mathbf{0.797} \pm 0.017$ |
| Myeloid | scGPT | $\mathbf{0.631} \pm 0.027$ | $0.338 \pm 0.012$ |
| | Geneformer | $0.619 \pm 0.014$ | $0.340 \pm 0.005$ |
| | scFoundation | $0.440 \pm 0.021$ | $0.251 \pm 0.004$ |
| | Cell2Sentence | $0.597 \pm 0.018$ | $0.577 \pm 0.018$ |
| | BioMod | $0.615 \pm 0.015$ | $\mathbf{0.645} \pm 0.016$ |

BMMC, Cell2Sentence achieved the higher overall score, with 0.677 compared with 0.662 for BioMod.

Table S3: **Five-run zero-shot integration metrics for BioMod.** Values are reported as mean ± s.d. across five independent zero-shot runs. Avg bio denotes biological conservation, Avg batch denotes batch mixing, and Avg score denotes the weighted overall score as defined in the Methods section. A compact comparison with Cell2Sentence is provided for metrics available in the corresponding evaluation results.

| Dataset or setting | NMI | ARI | Label ASW | Batch ASW | PCR batch | Graph conn. | Avg bio | Avg batch | Avg score |
|---|---|---|---|---|---|---|---|---|---|
| Kim Lung | $0.787 \pm 0.011$ | $0.649 \pm 0.019$ | $0.506 \pm 0.000$ | $0.975 \pm 0.000$ | $0.000 \pm 0.000$ | $0.988 \pm 0.001$ | $0.647 \pm 0.010$ | $0.982 \pm 0.000$ | $0.781 \pm 0.006$ |
| hPNC | $0.624 \pm 0.004$ | $0.388 \pm 0.004$ | $0.512 \pm 0.000$ | $0.958 \pm 0.000$ | $0.246 \pm 0.000$ | $0.844 \pm 0.012$ | $0.508 \pm 0.002$ | $0.901 \pm 0.006$ | $0.665 \pm 0.003$ |
| BMMC | $0.618 \pm 0.006$ | $0.377 \pm 0.024$ | $0.498 \pm 0.000$ | $0.972 \pm 0.000$ | $0.000 \pm 0.000$ | $0.843 \pm 0.004$ | $0.498 \pm 0.009$ | $0.907 \pm 0.002$ | $0.662 \pm 0.006$ |
| Kim Lung without malignant cells | $0.890 \pm 0.020$ | $0.885 \pm 0.041$ | $0.507 \pm 0.000$ | $0.977 \pm 0.000$ | $0.000 \pm 0.000$ | $0.996 \pm 0.001$ | $0.761 \pm 0.020$ | $0.986 \pm 0.000$ | $0.851 \pm 0.012$ |

*Compact comparison with Cell2Sentence.*

| Dataset or setting | Model | NMI | ARI | Label ASW | Avg bio | Avg batch | Avg score |
|---|---|---|---|---|---|---|---|
| Kim Lung | BioMod | 0.787 | 0.649 | 0.506 | 0.647 | 0.982 | **0.781** |
| Kim Lung | Cell2Sentence | 0.784 | 0.699 | 0.516 | 0.666 | 0.916 | 0.766 |
| hPNC | BioMod | 0.624 | 0.388 | 0.512 | 0.508 | 0.901 | **0.665** |
| hPNC | Cell2Sentence | 0.570 | 0.323 | 0.540 | 0.478 | 0.895 | 0.645 |
| BMMC | BioMod | 0.618 | 0.377 | 0.498 | 0.498 | 0.907 | 0.662 |
| BMMC | Cell2Sentence | 0.640 | 0.497 | 0.514 | 0.550 | 0.867 | **0.677** |
| Kim Lung without malignant cells | BioMod | 0.890 | 0.885 | 0.507 | 0.761 | 0.986 | **0.851** |
| Kim Lung without malignant cells | Cell2Sentence | 0.754 | 0.750 | 0.574 | 0.692 | 0.927 | 0.786 |

Table S4: **Comparison of model size and pretraining data scale across single-cell foundation models.** Model size and the number of pretraining cells are summarized for representative single-cell foundation models and BioMod.

| Model | Model size | Pretraining data scale |
|---|---|---|
| scBERT [18] | 6M | 1.1 million cells |
| scGPT [13] | 53M | 33.0 million cells |
| Geneformer [14] | 40M | 29.9 million cells |
| scFoundation [15] | 100M | 50.0 million cells |
| scLong [2] | 1B | 48.0 million cells |
| Regformer [19] | 50M | 26.87 million cells |
| Cell2Sentence [16] | 410M | 57.0 million cells |
| BioMod | **50M** | **73.0 million cells** |

#### S2.3 Perturbation prediction

We further evaluated BioMod on the Norman genetic perturbation prediction benchmark [20, 21] under four generalization settings: Seen 0/2, Seen 1/2, Seen 2/2, and Seen 0/1. For BioMod and all baseline models, mean squared error (MSE) and Pearson correlation were computed over the top 20 differentially expressed (DE) genes, consistent with the metrics reported in the shuffled-module comparison (Supplementary Table S11). Lower MSE indicates smaller prediction error, whereas higher Pearson correlation indicates stronger agreement between predicted and observed transcriptional responses.

BioMod achieved the lowest MSE in the Seen 1/2 setting, with a value of 0.180 (Supplementary Table S5). scLong [2] achieved the lowest MSE in the Seen 0/2 and Seen 0/1 settings, with values of 0.170 and 0.167, respectively, whereas scFoundation [15] achieved the lowest MSE of 0.119 in the Seen 2/2 setting. For Pearson correlation, BioMod achieved the highest values in the Seen 1/2, Seen 2/2 and Seen 0/1 settings, reaching 0.876, 0.953 and 0.876, respectively. scLong achieved the highest Pearson correlation of 0.893 in the Seen 0/2 setting. The ranking therefore changed across generalization settings and metrics, with BioMod showing its most consistent advantage in Pearson correlation.

Table S5: **Perturbation-response prediction performance on the Norman benchmark.** MSE and Pearson correlation are reported for four generalization settings: Seen 0/2, Seen 1/2, Seen 2/2, and Seen 0/1. For all models, both metrics were computed over the top 20 differentially expressed (DE) genes. Lower MSE and higher Pearson correlation indicate better performance. For baseline methods originally reported with uncertainty estimates, only point estimates are shown to keep the tabular comparison consistent. The best value for each metric is highlighted in bold.

| Model | 0/2 MSE ↓ | 0/2 Pearson ↑ | 1/2 MSE ↓ | 1/2 Pearson ↑ | 2/2 MSE ↓ | 2/2 Pearson ↑ | 0/1 MSE ↓ | 0/1 Pearson ↑ |
|---|---|---|---|---|---|---|---|---|
| GEARS | 0.218 | 0.862 | 0.221 | 0.819 | 0.130 | 0.855 | 0.191 | 0.561 |
| Geneformer | 0.185 | 0.860 | 0.237 | 0.799 | 0.195 | 0.805 | 0.170 | 0.576 |
| scGPT | 0.199 | 0.885 | 0.217 | 0.820 | 0.126 | 0.877 | 0.170 | 0.577 |
| scFoundation | 0.190 | 0.860 | 0.193 | 0.806 | **0.119** | 0.809 | 0.172 | 0.581 |
| UCE | 0.276 | 0.823 | 0.313 | 0.755 | 0.207 | 0.832 | 0.179 | 0.530 |
| scLong | **0.170** | **0.893** | 0.204 | 0.834 | 0.123 | 0.884 | **0.167** | 0.625 |
| BioMod | 0.175 | 0.871 | **0.180** | **0.876** | 0.122 | **0.953** | 0.171 | **0.876** |

To further characterize prediction errors in genetic perturbation-response prediction, we visualized gene-level error profiles across Norman perturbation settings. The heatmaps summarize the distribution of prediction errors across perturbation conditions and genes, allowing the identification of perturbation–gene regions with higher prediction difficulty. These visualizations complement the aggregate benchmark metrics by showing whether prediction errors are uniformly distributed or concentrated in specific perturbation–gene combinations.

#### S2.4 Drug-response prediction

BioMod achieved a five-seed mean Pearson correlation of $0.931 \pm 0.003$ for single-drug response prediction on the CCLE–GDSC benchmark [22–24] (Supplementary Table S6) and a five-seed mean AUROC of $0.681 \pm 0.004$ for drug-combination synergy prediction [25] (Supplementary Table S7).

#### S2.5 Module-level analyses of BioMod representations

Table S8 reports marker coverage for post-encoder token importance. A complementary input-level analysis and a random reference that accounts for marker membership in multiple modules are provided in Section S2.7.

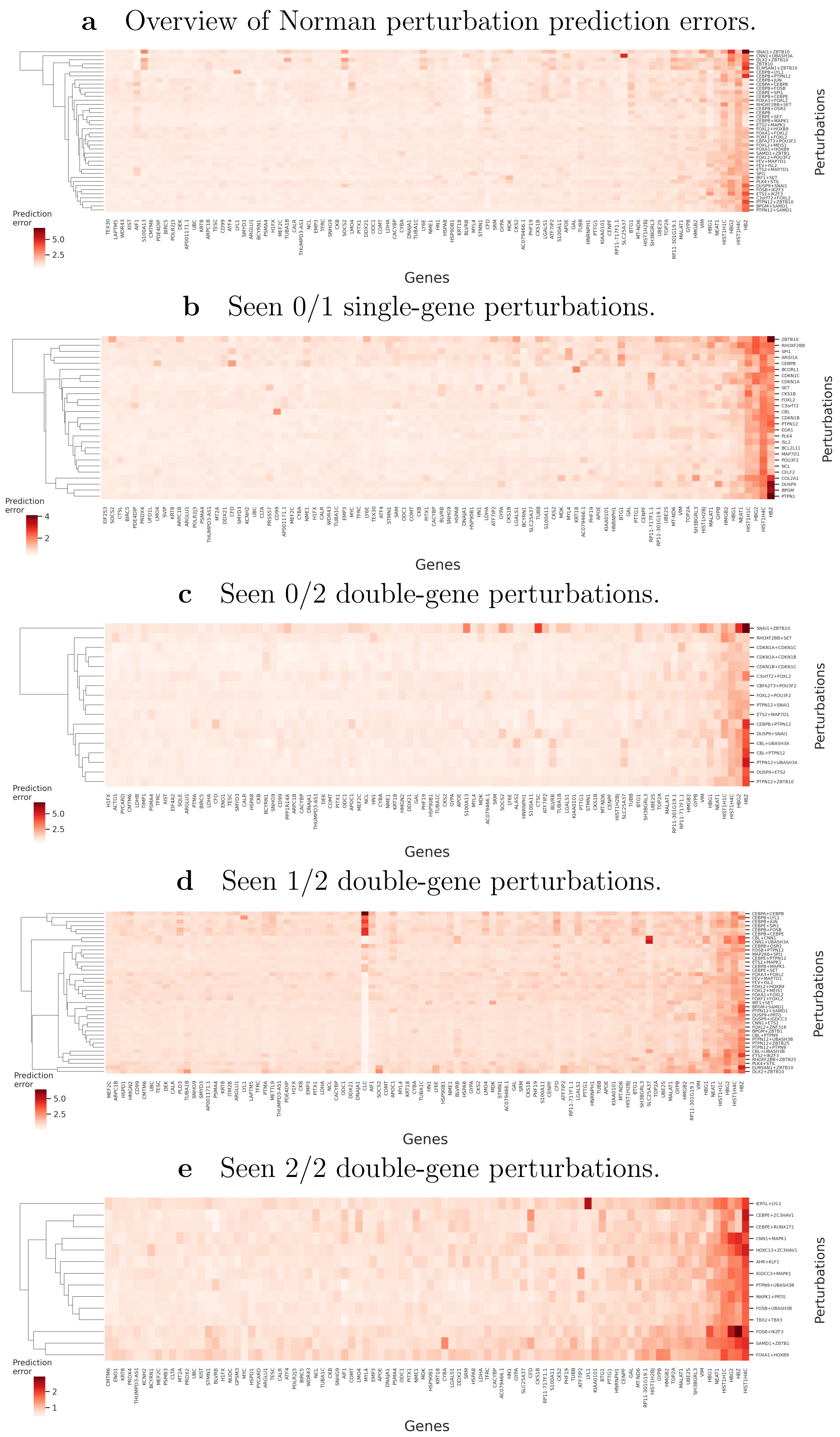

Figure S1: **Prediction-error profiles across Norman perturbation generalization settings. a,** Overview of gene-level prediction errors for Norman perturbations. **b–e,** Prediction errors for Seen 0/1 single-gene perturbations, Seen 0/2 double-gene perturbations, Seen 1/2 double-gene perturbations and Seen 2/2 double-gene perturbations, respectively. Rows represent perturbation conditions and columns represent genes. Color intensity denotes prediction error, with darker red indicating larger deviation between predicted and observed transcriptional responses. Hierarchical clustering was applied to perturbation conditions to reveal structured error patterns across perturbation classes.

Table S6: **Single-drug response prediction on the CCLE–GDSC benchmark test set.** Pearson correlation between predicted and observed $\ln(\mathrm{IC}_{50})$ values. Values are means across five seeds; standard deviations are shown for BioMod only.

| Model | Pearson |
|---|---|
| Linear | 0.746 |
| DeepCDR | 0.837 |
| Geneformer | 0.851 |
| scGPT | 0.841 |
| scFoundation | 0.867 |
| UCE | 0.837 |
| scLong | 0.878 |
| BioMod | **0.931** $\pm$ 0.003 |

Table S7: **Drug-combination synergy prediction.** AUROC for predicting whether a drug pair exhibits synergistic effects. Values are means across five seeds; standard deviations are shown for BioMod only.

| Model | AUROC |
|---|---|
| RandomForest | 0.533 |
| DeepDDS | 0.604 |
| Geneformer | 0.635 |
| scGPT | 0.616 |
| scFoundation | 0.593 |
| UCE | 0.603 |
| scLong | 0.652 |
| BioMod | **0.681** $\pm$ 0.004 |

Table S8: Canonical marker genes [26] contained in the top 5% highest-importance modules for each cell type in the Zheng68K dataset [8]. Importance was measured by zeroing individual module tokens after encoding and recording the decrease in the correct-class logit. Of 61 cell-type–marker entries, 59 mapped to at least one module; two unmapped entries were excluded from the rank summaries. Among the mapped entries, 46 were within the top 5% and 50 within the top 10% under post-encoder importance; this table lists the top-5% entries. The pre-encoder gene-set sensitivity results used in Fig. 6b of the main text are summarized in Supplementary Table S13, where 41 of 59 entries were within the top 5%.

| Cell type | Canonical marker genes in the top 5% importance modules |
|---|---|
| CD14+ Monocyte | CD14, LYZ, S100A8, S100A9, VCAN, FCN1, CD68 |
| CD19+ B | CD19, MS4A1, CD79A, CD79B, BANK1 |
| CD34+ | CD34, KIT, CRHBP |
| CD4+ T Helper2 | CD4, GATA3, IL7R, MAL |
| CD4+/CD25 T Reg | CD4, FOXP3, IL2RA, CTLA4 |
| CD4+/CD45RA+/CD25- Naive T | CD4, LEF1 |
| CD4+/CD45RO+ Memory | CD4, IL7R, S100A4, ANXA1, GPR183 |
| CD56+ NK | NCAM1, GNLY, KLRD1, KLRF1, GZMB |
| CD8+ Cytotoxic T | CD8A, CD8B, GZMB, NKG7 |
| CD8+/CD45RA+ Naive Cytotoxic | CD8A, CCR7, SELL, LEF1 |
| Dendritic | FCER1A, HLA-DRA, ITGAX |

#### S2.6 Ablation experiments

We report two complementary ablation analyses to evaluate the contribution of the module design in BioMod. First, the shuffled-module control examined whether biologically meaningful gene-to-module assignments were necessary for learning transferable cellular representations. Second, post-encoder module-family ablation quantified the contribution of Hallmark, Reactome, Hotspot and hdWGCNA representations [27–30] to cell-type classification.

For the shuffled-module experiment, the ablated model was pretrained from scratch using the same pretraining corpus, model architecture, optimization settings and downstream evaluation protocols as the complete model. The post-encoder module-family analysis did not involve model retraining and instead intervened directly on the module-token representations consumed by the fine-tuned classification head.

##### S2.6.1 Shuffled-module control

To determine whether the performance of BioMod depended on the biological composition of its modules rather than on gene grouping alone, we generated a shuffled-module control. Genes were randomly permuted and reassigned to modules according to the original module sizes. This procedure preserved the number of modules and the module-size distribution, while disrupting the predefined functional relationships among genes within each module. The shuffled-module model was subsequently pretrained from scratch using the randomized module definitions.

Table S9: Zero-shot batch-integration performance of the complete BioMod model and the shuffled-module control. The shuffled-module model preserved the original number of modules and module-size distribution but used randomized gene-to-module assignments.

| Dataset | Model | $NMI_{cluster/label}$ | $ARI_{cluster/label}$ | $ASW_{label}$ | $ASW_{label/batch}$ | $PCR_{batch}$ | Graph conn. | Avg bio | Avg batch | Avg score |
|---|---|---|---|---|---|---|---|---|---|---|
| Kim Lung | Complete | $0.787 \pm 0.011$ | $0.649 \pm 0.019$ | $0.506 \pm 0.000$ | $0.975 \pm 0.000$ | $0.000 \pm 0.000$ | $0.988 \pm 0.001$ | $0.647 \pm 0.010$ | $0.982 \pm 0.000$ | $0.781 \pm 0.006$ |
| Kim Lung | Shuffled | $0.767 \pm 0.014$ | $0.646 \pm 0.023$ | $0.482 \pm 0$ | $0.950 \pm 0$ | $0.000 \pm 0$ | $0.984 \pm 0.002$ | $0.632 \pm 0.012$ | $0.967 \pm 0$ | $0.766 \pm 0.007$ |
| hPNC | Complete | $0.624 \pm 0.004$ | $0.388 \pm 0.004$ | $0.512 \pm 0.000$ | $0.958 \pm 0.000$ | $0.246 \pm 0.000$ | $0.844 \pm 0.012$ | $0.508 \pm 0.002$ | $0.901 \pm 0.006$ | $0.665 \pm 0.003$ |
| hPNC | Shuffled | $0.242 \pm 0.008$ | $0.081 \pm 0.006$ | $0.487 \pm 0$ | $0.918 \pm 0$ | $0.225 \pm 0$ | $0.700 \pm 0.016$ | $0.270 \pm 0.004$ | $0.809 \pm 0.009$ | $0.486 \pm 0.005$ |
| BMMC | Complete | $0.618 \pm 0.006$ | $0.377 \pm 0.024$ | $0.498 \pm 0.000$ | $0.972 \pm 0.000$ | $0.000 \pm 0.000$ | $0.843 \pm 0.004$ | $0.498 \pm 0.009$ | $0.907 \pm 0.002$ | $0.662 \pm 0.006$ |
| BMMC | Shuffled | $0.584 \pm 0.009$ | $0.279 \pm 0.030$ | $0.492 \pm 0$ | $0.927 \pm 0$ | $0.000 \pm 0$ | $0.817 \pm 0.006$ | $0.452 \pm 0.012$ | $0.872 \pm 0.003$ | $0.620 \pm 0.008$ |
| Kim Lung without malignant cells | Complete | $0.890 \pm 0.020$ | $0.885 \pm 0.041$ | $0.507 \pm 0.000$ | $0.977 \pm 0.000$ | $0.000 \pm 0.000$ | $0.996 \pm 0.001$ | $0.761 \pm 0.020$ | $0.986 \pm 0.000$ | $0.851 \pm 0.012$ |
| Kim Lung without malignant cells | Shuffled | $0.725 \pm 0.026$ | $0.630 \pm 0.049$ | $0.484 \pm 0$ | $0.926 \pm 0$ | $0.000 \pm 0$ | $0.976 \pm 0.002$ | $0.613 \pm 0.023$ | $0.951 \pm 0$ | $0.748 \pm 0.014$ |

Shuffling gene-to-module assignments reduced the overall integration score in all four evaluation settings (Supplementary Table S9). Averaging the displayed scores across the four settings, the mean overall score decreased from approximately 0.740 to 0.655. The mean biological-conservation score decreased by approximately 0.112, compared with a decrease of approximately 0.044 in the mean batch-mixing score. The largest reduction was observed for hPNC, where the overall score decreased from 0.665 to 0.486.

Table S10: Cell-type annotation performance of the complete BioMod model and the shuffled-module control. $\Delta$ denotes Shuffled $-$ Complete.

| Dataset | Complete Acc. | Shuffled Acc. | $\Delta$ Acc. | Complete Macro-F1 | Shuffled Macro-F1 | $\Delta$ Macro-F1 |
|---|---|---|---|---|---|---|
| Zheng68K | $0.839 \pm 0.005$ | $0.809 \pm 0.007$ | $-0.030$ | $0.752 \pm 0.007$ | $0.714 \pm 0.010$ | $-0.038$ |
| Kim Lung | $0.986 \pm 0.002$ | $0.983 \pm 0.003$ | $-0.003$ | $0.962 \pm 0.005$ | $0.955 \pm 0.008$ | $-0.007$ |
| MS | $0.793 \pm 0.014$ | $0.636 \pm 0.020$ | $-0.157$ | $0.679 \pm 0.015$ | $0.538 \pm 0.022$ | $-0.141$ |
| Liver | $0.913 \pm 0.011$ | $0.893 \pm 0.014$ | $-0.020$ | $0.918 \pm 0.008$ | $0.893 \pm 0.012$ | $-0.025$ |
| Immune | $0.948 \pm 0.003$ | $0.893 \pm 0.006$ | $-0.055$ | $0.930 \pm 0.003$ | $0.861 \pm 0.006$ | $-0.069$ |
| Myeloid | $0.615 \pm 0.015$ | $0.627 \pm 0.019$ | $+0.012$ | $0.645 \pm 0.016$ | $0.339 \pm 0.025$ | $-0.306$ |
| hPancreas | $0.956 \pm 0.004$ | $0.950 \pm 0.006$ | $-0.006$ | $0.797 \pm 0.017$ | $0.783 \pm 0.021$ | $-0.014$ |
| Mean | 0.864 | 0.827 | $-0.037$ | 0.812 | 0.726 | $-0.086$ |

The shuffled-module model showed lower annotation accuracy on six of the seven datasets, with mean accuracy decreasing from 0.864 to 0.827 (Supplementary Table S10). Macro-F1 decreased on all seven datasets, with the mean decreasing from 0.812 to 0.726, a reduction of 0.086. The largest accuracy reduction was observed for MS, from 0.793 to 0.636. The largest macro-F1 reduction was observed for Myeloid, from 0.645 to 0.339. On Myeloid, accuracy increased from 0.615 to 0.627 despite the decrease in macro-F1, indicating that the two metrics reflected different aspects of annotation performance.

The effect of module shuffling on gene-perturbation prediction depended on the generalization setting and metric (Supplementary Table S11). MSE decreased from 0.175 to 0.160 in Seen 0/2 and from 0.171 to 0.162 in Seen 0/1, but increased from 0.180 to 0.189 in Seen 1/2 and from 0.122 to 0.149 in Seen 2/2. Pearson correlation decreased from 0.871 to 0.857 in Seen 0/2 and from 0.876 to 0.852 in Seen 1/2, while increasing slightly from 0.953 to 0.954 in Seen 2/2 and from 0.876 to 0.878 in Seen 0/1. In the drug-response ablation comparison reported in the same table, single-drug Pearson correlation decreased from 0.931 to 0.918, and drug-combination AUROC decreased from 0.681 to 0.628. Shuffling module assignments reduced performance in both drug-response tasks, whereas its effect on perturbation prediction varied across split definitions and metrics.

Table S11: Gene-perturbation and drug-response prediction performance of the complete BioMod model and the shuffled-module control. For gene-perturbation prediction, lower MSE and higher Pearson correlation indicate better performance.

| Task | Setting | Complete | Shuffled | Metric | Preferred direction |
|---|---|---|---|---|---|
| Gene perturbation | Seen 0/2 | 0.175 | 0.160 | Top-20 DE MSE | Lower |
| | | 0.871 | 0.857 | Top-20 DE Pearson | Higher |
| | Seen 1/2 | 0.180 | 0.189 | Top-20 DE MSE | Lower |
| | | 0.876 | 0.852 | Top-20 DE Pearson | Higher |
| | Seen 2/2 | 0.122 | 0.149 | Top-20 DE MSE | Lower |
| | | 0.953 | 0.954 | Top-20 DE Pearson | Higher |
| | Seen 0/1 | 0.171 | 0.162 | Top-20 DE MSE | Lower |
| | | 0.876 | 0.878 | Top-20 DE Pearson | Higher |
| Single-drug response | $0.931 \pm 0.003$ | $0.918 \pm 0.005$ | Pearson correlation | Higher | |
| Drug-combination response | $0.681 \pm 0.004$ | $0.628 \pm 0.007$ | AUROC | Higher | |

##### S2.6.2 Post-encoder module-family ablation

We performed post-encoder module-family ablation to quantify the contribution of Hallmark, Reactome, Hotspot and hdWGCNA representations [27–30] to cell-type classification on Zheng68K [8]. Unlike the shuffled-module experiment, this analysis did not involve model retraining.

Each test cell was first encoded by the fine-tuned backbone with the encoder masking ratio set to zero. For models using a non-CLS classification head, the encoded tokens were restored to their original module order. The resulting module-token representation was then modified before task-specific pooling and classification.

In the remove-one-family analysis, all module tokens belonging to the selected family were set to zero. In the retain-one-family and retain-two-family analyses, all tokens outside the selected family or family pair were set to zero. The modified module-token representation was subsequently passed through the original task-specific pooler and classifier. Therefore, the resulting changes in accuracy quantify the dependence of the fine-tuned classification model on the corresponding post-encoder module-family representations.

Among the single-family configurations, retaining only Reactome modules produced the highest classification accuracy (0.713), compared with 0.658, 0.676 and 0.643 for Hallmark, Hotspot and hdWGCNA, respectively (Supplementary Table S12). Removing Reactome produced the largest decrease among the remove-one-family experiments, reducing accuracy from 0.839 to 0.764, a change of −0.075. Removing Hallmark, Hotspot or hdWGCNA resulted in smaller reductions, with accuracies of 0.802, 0.819 and 0.821 and corresponding changes of −0.037, −0.020 and −0.018, respectively.

Among the two-family configurations, Reactome combined with Hotspot achieved the highest accuracy (0.788), followed by Reactome with hdWGCNA (0.759) and Hallmark with Reactome (0.756). Each Reactome-containing pair outperformed the family pairs that excluded Reactome. However, none of the single-family or two-family configurations matched the accuracy of the complete model.

In this Zheng68K analysis, Reactome was the largest individual contributor to cell-type classification, and the full four-family representation gave the highest accuracy. Because

Table S12: Post-encoder module-family ablation on the Zheng68K cell-type annotation task. The complete model used all four module families. The baseline accuracy was recomputed using the unmodified module-token representation in the same post-encoder evaluation pipeline. Δ denotes the change in accuracy relative to the complete model.

| Ablation setting | Retained or removed family | Accuracy | Δ accuracy |
|---|---|---|---|
| Complete model | All four families | 0.839 | – |
| Retain one | Hallmark only | 0.658 | -0.181 |
| Retain one | Reactome only | 0.713 | -0.126 |
| Retain one | Hotspot only | 0.676 | -0.163 |
| Retain one | hdWGCNA only | 0.643 | -0.196 |
| Remove one | Remove Hallmark | 0.802 | -0.037 |
| Remove one | Remove Reactome | 0.764 | -0.075 |
| Remove one | Remove Hotspot | 0.819 | -0.020 |
| Remove one | Remove hdWGCNA | 0.821 | -0.018 |
| Retain two | Hallmark + Reactome | 0.756 | -0.083 |
| Retain two | Hallmark + Hotspot | 0.706 | -0.133 |
| Retain two | Hallmark + hdWGCNA | 0.679 | -0.160 |
| Retain two | Reactome + Hotspot | 0.788 | -0.051 |
| Retain two | Reactome + hdWGCNA | 0.759 | -0.080 |
| Retain two | Hotspot + hdWGCNA | 0.682 | -0.157 |

the intervention was applied after encoder computation, these results should be interpreted as the dependence of the fine-tuned classification head on different module-family representations, rather than as a comparison of family-specific pretraining effectiveness.

#### S2.7 Pre-encoder gene-set sensitivity

To assess how module rankings depended on the intervention site, we compared pre-encoder gene-set sensitivity with post-encoder token importance on Zheng68K. The comparison used the same fine-tuned checkpoint and fixed stratified test split of 13,690 cells, comprising 11 cell types and 660 modules. For each module, we set its constituent genes' expression values to zero, reran the full encoder and classifier, and averaged the decrease in the correct-class logit within each cell type. Post-encoder importance instead measured the effect of zeroing an individual encoded module token.

The two rankings showed moderate within-cell-type concordance (Supplementary Fig. S2 and Table S13). The median Spearman correlation was 0.463 across cell types, with a range of 0.287–0.575. The median overlap between the two top-50 module sets was 40%, ranging from 16% to 70%. After averaging importance equally across the 11 cell types, the correlation across 660 modules was 0.336. This module-mean statistic differs from the correlation of 0.130 obtained by pooling all 7,260 cell-type–module entries without averaging.

Canonical marker coverage was retained under input-level perturbation. Of 61 cell-type–marker entries, 59 mapped to at least one module. For each mapped entry, we recorded the highest rank among modules containing that marker in the corresponding cell-type ranking. Pre-encoder sensitivity placed 41 of 59 entries (69.5%) within the top 5% of modules, compared with 46 of 59 (78.0%) for post-encoder importance. Their median rank percentiles were 0.91% and 1.97%, respectively, using $100(r-1)/660$ for a best-module rank of $r$.

To account for markers occurring in multiple modules, we drew 33 of the 660 modules uniformly without replacement for each cell type and counted marker entries represented in at least one selected module. The original marker-to-module memberships were retained. Across 20,000 random draws (seed 20260912), the expected total was 15.135 of the 59 mapped entries (25.7%). No random draw reached either observed count, giving a one-sided empirical $P = 1/20{,}001 \approx 5.0 \times 10^{-5}$ for each ranking under the finite-sample correction $(b+1)/(B+1)$, where $b$ is the number of random counts at least as large as the observed count and $B = 20{,}000$. Thus, selecting the top 5% of modules does not imply a 5% random expectation for marker coverage. This reference accounts for marker membership in multiple modules but does not match modules by size or measure uncertainty across model-training runs.

Pre-encoder sensitivity was associated with module size: the Spearman correlation between module gene count and mean importance was 0.378, compared with −0.102 for post-encoder importance. The 50 modules with the highest mean pre-encoder importance contained 173.3 genes on average, compared with 105.5 across all modules. Among genes represented in the module collection, 70.1% occurred in more than one module, with a mean of 5.21 module memberships per gene. Consequently, zeroing one module's genes also changes the inputs to overlapping modules and propagates through the encoder. These results describe sensitivity to overlapping gene sets, not independent causal contributions of individual pathways. The CD4+ T Helper2 and CD34+ rankings are descriptive because their test sets contained only 19 and 49 cells, respectively. PanglaoDB analysis returned 1,609 significant results among 5,874 tests at adjusted $P < 0.05$, but the best-matching database labels often differed from the target cell types; these matches were not treated as direct validation of cell identity. Pre-encoder sensitivity therefore complements, rather than replaces, the post-encoder ranked-token ablation evidence.

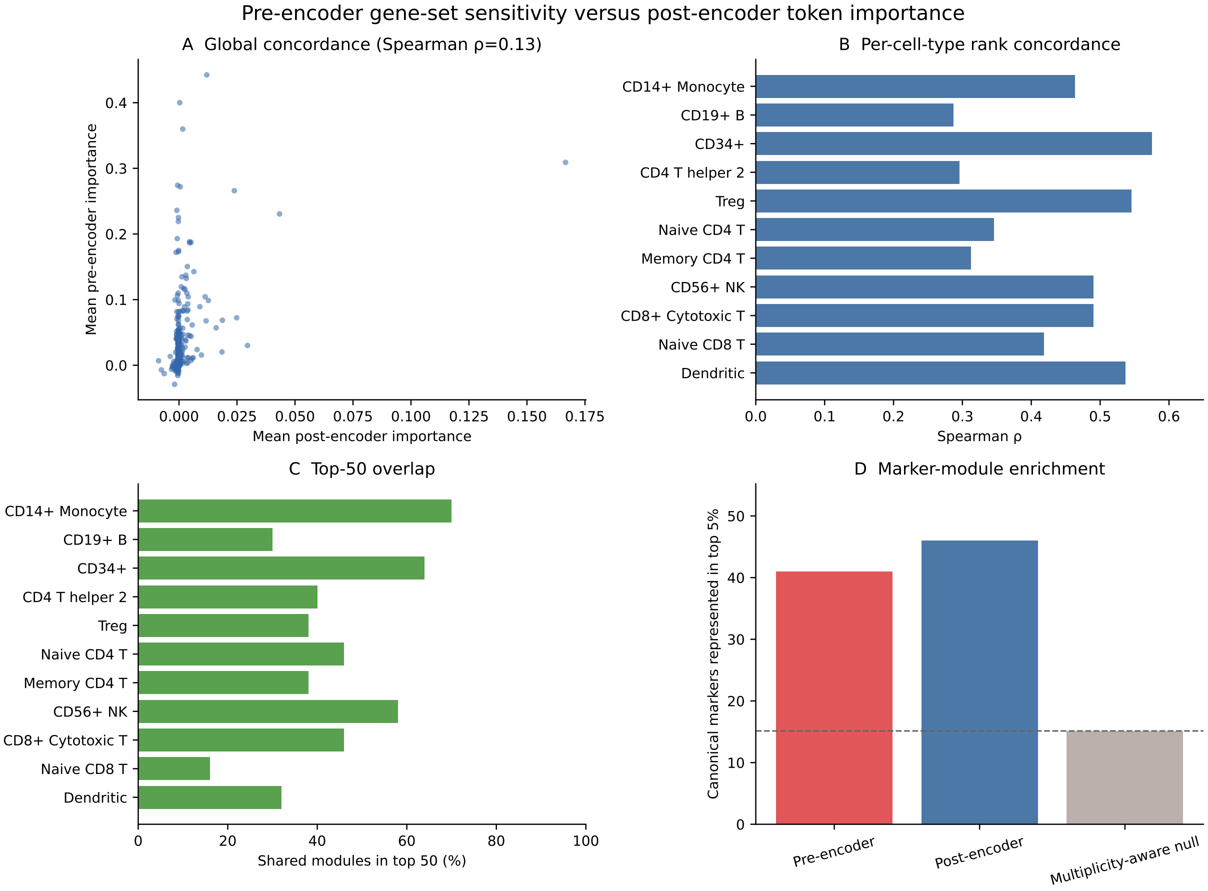

Figure S2: **Comparison of pre-encoder gene-set sensitivity and post-encoder token importance.** **a**, Mean importance across 11 cell types for each of 660 modules; the Spearman correlation is computed from these 660 pairs of module means. **b**, Within-cell-type Spearman correlation across modules. **c**, Percentage of modules shared by the respective top-50 sets. **d**, Number of mapped cell-type–marker entries represented in the top 5% of modules under each ranking. The gray bar and dashed line show the mean count from 20,000 random draws of 33 modules per cell type, retaining marker-to-module memberships. The analysis used 13,690 test cells; per-class sample sizes are listed in Supplementary Table S13. The panels summarize the stored importance matrices and are not averages across training runs.

Table S13: **Cell-type-specific agreement between pre-encoder and post-encoder module rankings.** $n$ denotes test cells; $\rho$ is Spearman correlation across 660 modules; overlap is the percentage shared by the two top-50 sets. The final two columns report mapped cell-type–marker entries represented in the top 5% of modules, with the number of mapped entries as denominator. The same gene may contribute an entry for more than one cell type. Rankings for the two smallest classes should be interpreted descriptively.

| Cell type | $n$ | $\rho$ | Top-50 overlap (%) | Pre markers | Post markers |
|---|---|---|---|---|---|
| CD14+ Monocyte | 570 | 0.463 | 70 | 7/7 | 7/7 |
| CD19+ B | 1,175 | 0.287 | 30 | 5/6 | 5/6 |
| CD34+ | 49 | 0.575 | 64 | 3/4 | 3/4 |
| CD4+ T Helper2 | 19 | 0.296 | 40 | 3/5 | 4/5 |
| CD4+/CD25 T Reg | 1,237 | 0.546 | 38 | 3/5 | 4/5 |
| CD4+/CD45RA+/CD25- Naive T | 374 | 0.346 | 46 | 2/5 | 2/5 |
| CD4+/CD45RO+ Memory | 612 | 0.313 | 38 | 4/5 | 5/5 |
| CD56+ NK | 1,755 | 0.491 | 58 | 5/6 | 5/6 |
| CD8+ Cytotoxic T | 4,151 | 0.491 | 46 | 6/6 | 4/6 |
| CD8+/CD45RA+ Naive Cytotoxic | 3,329 | 0.419 | 16 | 1/5 | 4/5 |
| Dendritic | 419 | 0.537 | 32 | 2/5 | 3/5 |

#### S2.8 Inference efficiency

The following tables report the representation-forward measurements plotted in the main-text inference-efficiency panel. A fixed cohort of 10,000 Zheng68K cells was selected with seed 42. The guarded pool-affine BioMod implementation was measured on the first 512 cells of this cohort, whereas each comparator was measured on all 10,000 cells. Models retained their respective input adapters and sequence lengths. Measurements ran sequentially on the same NVIDIA H100 80-GB GPU with BF16 representation weights and autocast. For each batch-size setting, 30 warm-up batches preceded five technical timing repeats. The synchronized forward-time boundary included learned input processing, the representation backbone, normalization and cell-level pooling; it excluded CPU preprocessing, host-to-device transfer and unused reconstruction decoders or output heads. The unequal cohort sizes make the cross-model comparisons descriptive rather than matched full-cohort acceleration estimates.

Table S14: **Representation-forward throughput across batch sizes.** Values are cells per second (mean ± sample standard deviation across five technical repeats). BioMod used 512 cells per repeat; each comparator used 10,000. OOM indicates CUDA out-of-memory failure, for which no throughput value was assigned.

| Model | Batch 1 | Batch 16 | Batch 64 | Batch 256 |
|---|---|---|---|---|
| BioMod | 364.07 ± 4.72 | 1391.14 ± 3.20 | 1729.36 ± 20.50 | 1761.07 ± 22.85 |
| scGPT | 269.97 ± 1.09 | 1137.48 ± 0.41 | 1311.79 ± 0.23 | 1231.04 ± 0.34 |
| Geneformer | 211.27 ± 0.94 | 571.90 ± 0.39 | 386.66 ± 0.08 | OOM |
| scFoundation | 224.72 ± 1.52 | 596.78 ± 1.47 | 673.46 ± 1.50 | 532.21 ± 0.23 |
| scLong | 7.90664 ± 0.00052 | 8.78530 ± 0.00028 | 8.70330 ± 0.00034 | OOM |
| Cell2Sentence | 73.34 ± 0.29 | 157.53 ± 0.05 | 163.59 ± 0.02 | 163.33 ± 0.14 |

Table S15: **Single-cell representation-forward latency at batch size 1.** Values are median [first quartile, third quartile] in milliseconds, pooled across five technical repeats. BioMod contributed 2,560 timed forwards from 512 cells per repeat; each comparator contributed 50,000 from 10,000 cells per repeat.

| Model | Median [Q1, Q3] (ms) | Timed forwards |
|---|---|---|
| BioMod | 2.70 [2.68, 2.71] | 2,560 |
| scGPT | 3.69 [3.65, 3.72] | 50,000 |
| Geneformer | 4.68 [4.65, 4.71] | 50,000 |
| scFoundation | 4.42 [4.36, 4.46] | 50,000 |
| scLong | 126.41 [126.39, 126.43] | 50,000 |
| Cell2Sentence | 13.57 [13.36, 13.74] | 50,000 |

[5] Jingcheng Du, Peilin Jia, Yulin Dai, Cui Tao, Zhongming Zhao, and Degui Zhi. Gene2vec: distributed representation of genes based on co-expression. *BMC Genomics*, 20(Suppl 1):82, 2019.

[6] Ziyang Wu, Tianjiao Ding, Yifu Lu, Druv Pai, Jingyuan Zhang, Weida Wang, Yaodong Yu, Yi Ma, and Benjamin D. Haeffele. Token statistics transformer: Linear-time attention via variational rate reduction. *arXiv preprint arXiv:2412.17810*, 2024.

[7] Kaiming He, Xinlei Chen, Saining Xie, Yanghao Li, Piotr Dollár, and Ross Girshick. Masked autoencoders are scalable vision learners. In *Proceedings of the IEEE/CVF Conference on Computer Vision and Pattern Recognition (CVPR)*, pages 16000–16009, 2022.

[8] Grace X. Y. Zheng, Jessica M. Terry, Phillip Belgrader, Paul Ryvkin, Zachary W. Bent, Ryan Wilson, Solongo B. Ziraldo, Tobias D. Wheeler, Geoff P. McDermott, Junjie Zhu, Mark T. Gregory, Joe Shuga, Luz Montesclaros, Jason G. Underwood, Donald A. Masquelier, Stefanie Y. Nishimura, Michael Schnall-Levin, Paul W. Wyatt, Christopher M. Hindson, Rajiv Bharadwaj, Alexander Wong, Kevin D. Ness, Lan W. Beppu, H. Joachim Deeg, Christopher McFarland, Keith R. Loeb, William J. Valente, Nolan G. Ericson, Emily A. Stevens, Jerald P. Radich, Tarjei S. Mikkelsen, Benjamin J. Hindson, and Jason H. Bielas. Massively parallel digital transcriptional profiling of single cells. *Nature Communications*, 8(1):14049, 2017.

[9] Nayoung Kim, Hong Kwan Kim, Kyungjong Lee, Yourae Hong, Jhingook H. Cho, Jung Won Choi, Jung-Il Lee, Yeon-Lim Suh, Bo Mi Ku, Hye Hyeon Eum, Soyean

Choi, Yoon-La Choi, Je-Gun Joung, Woong-Yang Park, Hye Ryun Jung, Jong-Mu Sun, Se-Hoon Lee, Jin Seok Ahn, Keunchil Park, Myung-Ju Ahn, and Hae-Ock Lee. Single-cell RNA sequencing demonstrates the molecular and cellular reprogramming of metastatic lung adenocarcinoma. *Nature Communications*, 11(1):2285, 2020.

[22] Jordi Barretina, Giordano Caponigro, Nicolas Stransky, Kavitha Venkatesan, Adam A. Margolin, Sungjoon Kim, Christopher J. Wilson, Joseph Lehár, Gregory V. Kryukov, Dmitriy Sonkin, Anupama Reddy, Manway Liu, Lauren Murray, Michael F. Berger, John E. Monahan, Paula Morais, Jodi Meltzer, Adam Korejwa, Judit Jané-Valbuena, Felipa A. Mapa, Joseph Thibault, Eva Bric-Furlong, Pichai Raman, Aaron Shipway, Ingo H. Engels, Jill Cheng, Guoying K. Yu, Jianjun Yu, Peter Jr Aspesi, Melanie de Silva, Kalpana Jagtap, Michael D. Jones, Li Wang, Charles Hatton, Emanuele Palescandolo, Supriya Gupta, Scott Mahan, Carrie Sougnez, Robert C. Onofrio, Ted Liefeld, Laura MacConaill, Wendy Winckler, Michael Reich, Nanxin Li, Jill P. Mesirov, Stacey B. Gabriel, Gad Getz, Kristin Ardlie, Vivien Chan, Vic E. Myer, Barbara L. Weber, Jeff Porter, Markus Warmuth, Peter Finan, Jennifer L. Harris, Matthew Meyerson, Todd R. Golub, Michael P. Morrissey, William R. Sellers, Robert Schlegel, and Levi A. Garraway. The cancer cell line encyclopedia enables predictive modelling of anticancer drug sensitivity. *Nature*, 483(7391):603–607, 2012.

[23] Wanjuan Yang, Jorge Soares, Patricia Greninger, Elena J. Edelman, Howard Lightfoot, Simon Forbes, Nidhi Bindal, Dave Beare, James A. Smith, I. Richard Thompson, Sridhar Ramaswamy, P. Andrew Futreal, Daniel A. Haber, Michael R. Stratton, Cyril Benes, Ultan McDermott, and Mathew J. Garnett. Genomics of drug sensitivity in cancer (GDSC): a resource for therapeutic biomarker discovery in cancer cells. *Nucleic Acids Research*, 41(D1):D955–D961, 2013.

[24] Qiao Liu, Zhiqiang Hu, Rui Jiang, and Mu Zhou. DeepCDR: a hybrid graph convolutional network for predicting cancer drug response. *Bioinformatics*, 36(Supplement_2):i911–i918, 2020.

[25] Jinxian Wang, Xuejun Liu, Siyuan Shen, Lei Deng, and Hui Liu. DeepDDS: deep graph neural network with attention mechanism to predict synergistic drug combinations. *Briefings in Bioinformatics*, 23(1):bbab390, 2022.

[26] Oscar Franzén, Li-Ming Gan, and Johan L. M. Björkegren. PanglaoDB: a web server for exploration of mouse and human single-cell RNA sequencing data. *Database*, 2019:baz046, 2019.

[27] Arthur Liberzon, Chet Birger, Helga Thorvaldsdóttir, Mahmoud Ghandi, Jill P. Mesirov, and Pablo Tamayo. The molecular signatures database (MSigDB) hallmark gene set collection. *Cell Systems*, 1(6):417–425, 2015.

[28] Marc Gillespie, Bijay Jassal, Ralf Stephan, Marija Milacic, Karen Rothfels, Amanda Senff-Ribeiro, Johannes Griss, Cristoffer Sevilla, Lisa Matthews, Chuqiao Gong, Chuqiao Deng, Thawfeek Varusai, Eliot Ragueneau, Yusuf Haider, Bruce May, Veronica Shamovsky, Joel Weiser, Timothy Brunson, Nima Sanati, Linda Beckman, Xiaoming Shao, Antonio Fabregat, Konstantinos Sidiropoulos, Juan Murillo, Guilherme Viteri, Justin Cook, Solomon Shorser, Gary Bader, Emek Demir, Chris Sander, Robin Haw, Guanming Wu, Lincoln Stein, Henning Hermjakob, and Peter D'Eustachio. The Reactome pathway knowledgebase 2022. *Nucleic Acids Research*, 50(D1):D687–D692, 2022.

[29] David DeTomaso and Nir Yosef. Hotspot identifies informative gene modules across modalities of single-cell genomics. *Cell Systems*, 12(5):446–456.e9, 2021.

[30] Samuel Morabito, Fairlie Reese, Negin Rahimzadeh, Emily Miyoshi, and Vivek Swarup. hdWGCNA identifies co-expression networks in high-dimensional transcriptomics data. *Cell Reports Methods*, 3(6):100498, 2023.
